# Apaf-1 Pyroptosome Senses Mitochondrial Permeability Transition

**DOI:** 10.1101/2020.01.27.921122

**Authors:** Wanfeng Xu, Yuan Che, Quan Zhang, Hai Huang, Chujie Ding, Yun Wang, Guangji Wang, Lijuan Cao, Haiping Hao

## Abstract

Caspase-4 directly senses and is activated by cytosolic LPS in conditions of pathogen infection. It is unclear whether and how caspase-4 detects host derived factors for triggering pyroptosis. Here we show that mitochondrial permeability transition (MPT) promotes the assembly of a protein complex comprised of Apaf-1 and caspase-4 (caspase-11 in mice), defined herein as pyroptosome, for the execution of facilitated pyroptosis. MPT induced by bile acids and calcium overload, and specifically by an adenine nucleotide translocator 1 (ANT1) activator, triggered pyroptosome assembly. Different from the direct cleavage of GSDMD by LPS-activated caspase-4, caspase-4 activated in the Apaf-1 pyroptosome proceeds to cleave caspase-3 and thereby gasdermin E (GSDME) to induce pyroptosis. Caspase-11 initiated and GSDME executed pyroptosis underlies cholesteric liver failure. These findings identify Apaf-1 pyroptosome as a pivotal machinery for cells sensing MPT signals and may shed lights on understanding how cells execute pyroptosis under sterile conditions.

**Highlights:**

- Bile acids trigger caspase-4/11 and GSDME dependent pyroptosis
- Caspase-4/11 is a general sensor of mitochondrial permeability transition (MPT)
- MPT drives Apaf-1/capase-4 pryoptosome assembly
- Caspase-11 and GSDME mediated pyroptosis underlies cholesteric liver damage

**eTOC Blurb:** Persistent mitochondrial permeability transition elicited by bile acids, calcium overload and specifically ANT1 activators drives assembly of Apaf-1-capase-4/11 pyroptosome triggering GSDME dependent pryroptosis.

## INTRODUCTION

Pyroptosis is typically featured with pore formation on plasma membrane, cell lysis and swelling, and is mediated by inflammatory caspases (Jorgensen and Miao, 2015). Both caspase-1 related canonical inflammasome(Miao et al., 2010) and caspase4/5/11 in non-canonical inflammasome have been found to play key roles in pyroptosis(Kayagaki et al., 2011). Caspase-1 is activated on the protein complex platform called inflammasome, among of which various pattern recognition receptors sense pathogen associated molecular patterns (PAMPs) or host derived danger associated molecular patterns (DAMPs)(Man and Kanneganti, 2015). Activated caspase-1 cleaves IL-1β or IL-18 to trigger inflammation, and gasdermin D (GSDMD) to induce pyroptosis(Bergsbaken et al., 2009; Russo et al., 2016; Sborgi et al., 2016; Shi et al., 2015; Taabazuing et al., 2017). In contrast to the extensively studied mechanisms about activation of canonical inflammasomes, the understanding of how caspase-4, previously defined as noncanonical inflammasome, is activated is only emerging. Caspase-4/11 can directly detect cytoplasmic LPS. Caspase-4/11 CARD domain interacts with a carbohydrate chain of LPS called lipid A, leading to caspase-11 auto-cleavage and subsequently pyroptotic executioners activation, providing a mechanistic rationale for understanding why caspase-4/11 plays pivotal roles in pyroptotic cell death implicated in infectious diseases(Hagar et al., 2013; Shi et al., 2014).

Notably, in addition to microbial infections, pyroptosis has been widely observed in sterile conditions implicated in many types of diseases. The mechanism and signal pathways underlying pyroptosis under sterile conditions have been largely attributed to the activation of canonical inflammasome and caspase-1. It is not a surprise considering that canonical inflammasome is able to sense diverse DAMPs and endogenous bioactive metabolites such as fatty acids, amino acids and bile acids(Hao et al., 2017; Levy et al., 2015; Pan et al., 2018). Of interest, caspase-4/11 can also detect endogenous oxidized phospholipids facilitating IL-1β release but not pyroptosis(Zanoni et al., 2016). It is important to note that caspase-4/11 is widely distributed in diverse organs/tissues where its pathophysiological significance remains largely elusive. A critical question is that whether and how caspase-4/11 detects other host derived endogenous factors and thereby triggering pyroptosis implicated in noninfective diseases.

Mitochondrion is a central hub in sensing various stress stimuli and thereby controlling cell fates including cell death, differentiation, proliferation and immunological responses (Kroemer et al., 2007; Mehta et al., 2017). Generally, moderate stress may lead to mitochondrial outer membrane permeabilization (MOMP) to trigger apoptotic cell death via eliciting pro-apoptotic factors activation. Mitochondria also regulate other forms of cell deaths, including necroptosis and ferroptosis(Bock and Tait, 2019). An increasing amount of evidences indicates that the mitochondrial permeability transition (MPT) acts as a key nodal in regulating necrosis(Nakagawa et al., 2005). MPT is induced by the persistent opening of the mitochondrial permeability transition pore (MPTP), usually activated by Ca^2+^ together with phosphate and reactive oxygen species (ROS)(Hurst et al., 2017). MPT induced cell death has been widely supposed to be a kind of regulated necrosis; however, it remains largely unclear about what is the regulated necrosis and how cells sense MPT for culminating cell death.

Here, we provide direct evidence supporting that caspase-4/11 represents a general sensor of MPT induced by bile acids, calcium overload and specifically by activators of adenine nucleotide translocator (ANT). MPT activates Apaf-1 that surprisingly recruit and activate caspase-4/11, but not caspase-9. Caspase-4/11 activated in the Apaf-1 pyroptosome cleaves caspase-3 that further cleaves gasermine E (GSDME) for eliciting pyroptosis. These findings provide a causal link of capase-4/11 activation mediated pyroptosis implicated in diverse sterile diseases and support that mitochondrion is also a central hub in regulating pyroptosis, in addition to previously well-established other types of cell death.

## RESULTS

### Bile Acids Trigger Caspase-4/11 Dependent Pyroptosis

We have recently identified bile acids as a class of distinctive DAMPs which can activate both signal 1 and 2 of NLRP3 inflammasome in mature macrophages but not in monocytes(Hao et al., 2017). Bile acids are known for their cytotoxicity but without clear molecular events(Katona et al., 2009). Because inflammasome activation is connected with pyroptosis, we asked whether bile acids may also trigger pyroptotic cell death. Both deoxycholic acid (DCA) and chenodeoxycholic acid (CDCA) activated caspase-11 (caspase-4 in THP-1 cells) and promoted the release of HMGB1, IL-1α and cell death in mouse bone marrow derived macrophages (BMDMs) (**Figures 1A-F**) as well as in THP-1 cells (**Figures S1A-D**). Interestingly, bile acid concomitantly activated caspase-11 and −3, which are respectively key executers of pyroptosis and apoptosis (**Figure 1F**). Canonical inflammasome represented by caspase-1 activation and IL-1β secretion was not influenced by bile acids treatment without LPS priming in BMDMs (**Figure 1D and 1F**), which agrees with our previous findings that bile acids alone without LPS priming were hard to activate Nlrp3 inflammasome in BMDMs, in spite of caspase-1 activation in THP-1 cells(Hao et al., 2017) (**Figure S1A**). We validated that bile acids induced cell death was typical pyroptosis as evidenced from morphological assessment and propidium iodide (PI) positive staining (**Figures 1G and 1H, Figure S1E**). Tauro-conjugates of DCA and CDCA (TDCA and TCDCA), which were capable of activating signal 2 of NLRP3 inflammasome(Hao et al., 2017), could not activate caspase-4/11 and were unable to trigger pyroptosis (**Figures S1C-I**), indicating that bile acids active pyroptotic signals via a mechanism distinct to that for activation of canonical inflammasome.

**Figure 1.**
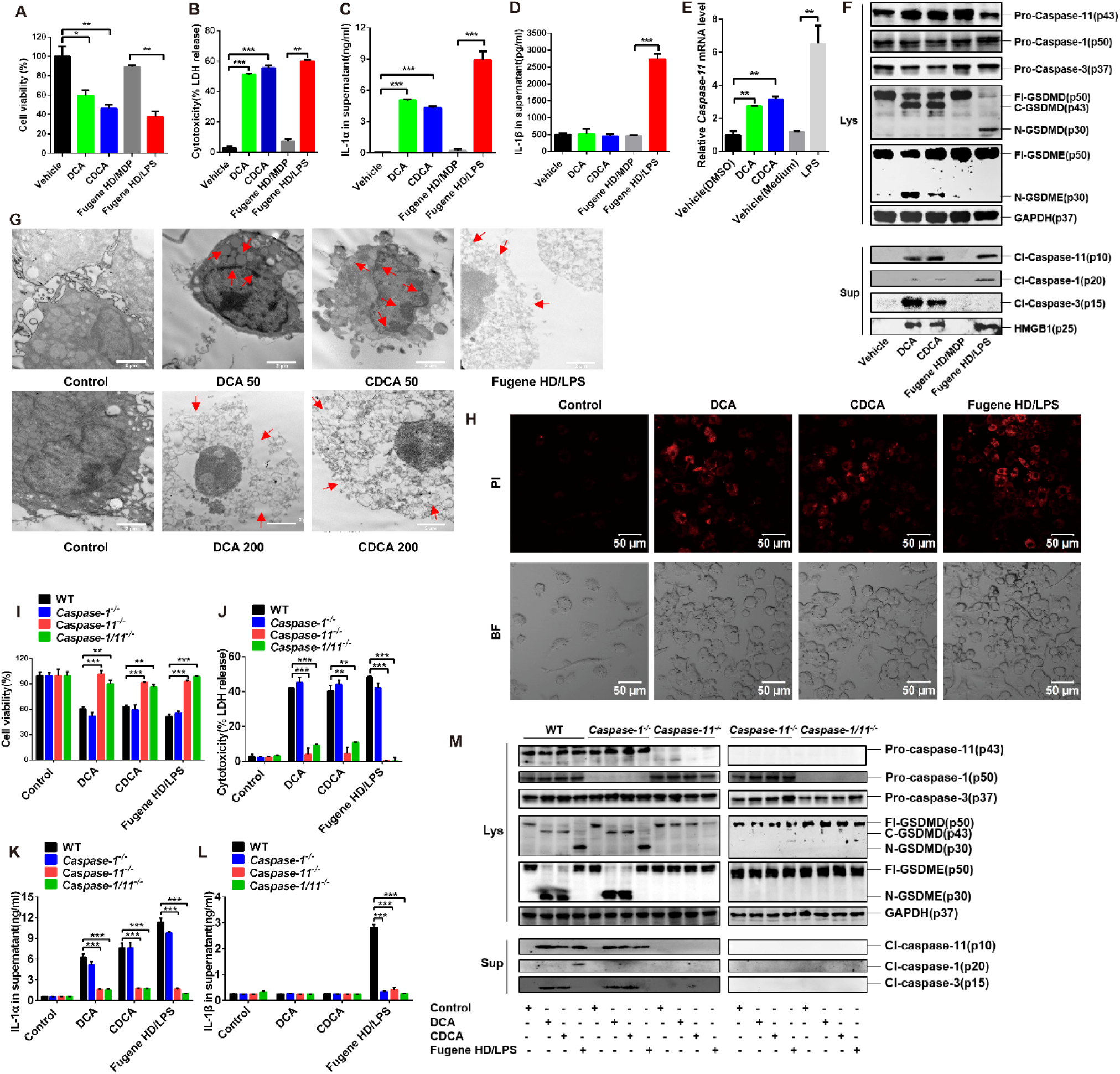
Bile Acids Activate Caspase-11 Mediated Pyroptosis in BMDMs. **(A-D)** CCK8 assay for cell viability **(A)**, LDH release for cytotoxicity **(B)** and IL-1α/β secretion **(C and D)** of BMDMs treated with 200 μM bile acids (DCA, CDCA) for 4 hrs, or for LPS-induced pyroptosis, BMDMs were primed with 1ug/mL LPS for 6 hrs followed by transfection with 5 μg/mL LPS by 0.3% (v/v) Fugene HD for another 16 hrs; equimolar muramyl dipeptide (MDP) transfected by Fugene HD was set as a negative control. **(E)** *Caspase-11* mRNA levels in BMDMs treated with 50 ng/ml LPS or 200 μ BAs for 4 hrs. **(F)** Cell lysates (Lys) or supernatant (Sup) of BMDMs were collected for immunoblot analysis of caspase-1/11/3, GSDME/D and HMGB1. Experimental design as in (A-D). **(G)** Representative images of electron microscopy analysis of apoptotic or pyroptotic morphology changes in BMDMs. Cells were stimulated with 50 μM bile acid for 16 hrs or 200 μM bile acids for 4 hrs; for LPS group, LPS-primed cells were transfected with 5 μg/mL LPS by 0.3% Fugene HD for 16 hrs. Scale bars, 2 μm. **(H)** Representative confocal microscopy analysis of PI positive cells, scale bars, 50 μm. Experimental design as in (A-D). **(I-L)** CCK8 assay for cell viability **(I)**, DH release for cytotoxicity **(J)** and IL-1α β secretion **(K and L)** of BMDMs isolated from WT, *Caspase-1^-/-^*, *Caspase-11^-/-^*, *Caspase-1/11^-/-^* mice. Experimental design as in (A-D). **(M)** Immunoblot analysis of caspase-1/11/3, GSDME/D, HMGB1 and GAPDH in cell lysates (Lys) and supernatant (Sup) of BMDMs isolated from WT, *Caspase-1^-/-^*, *Caspase-11^-/-^*, *Caspase-1/11^-/-^* mice. Experimental design as in (A-D). GAPDH was used as internal standard/loading control in qPCR/immunoblot analyses. Bar graphs expressed as mean ± SEM (n=3). \*\*\**p*<0.001, ** *p* < 0.01, * *p* < 0.05 compared to control unless indicated otherwise in graphs. See also **Figure S1**.

Bile acids induced cell death, LDH release and IL-1α secretion were dramatically reduced in BMDMs isolated from *Caspase-11^-/-^* mice in comparison with that from wild-type (WT) mice. In contrast, knockout of caspase-1 exerted little influence on bile acid induced cell death and IL-1α secretion, confirming that the featured cell death triggered by bile acid is caspase-11 mediated pyroptosis (**Figures 1I-L**). Cytotoxicity of bile acids at concentrations higher than 400 μM might be partially related to their detergent effects (Benedetti et al., 1997). We further demonstrated that equivalent amount of SDS to that of 200 μM bile acid was unable to activate caspases, excluding the possibility of detergent effects being involved in caspase activation by bile acids (**Figures S1J-M**). Neither RIP1 inhibitor nor MLKL inhibitor blocked bile acids induced cell death, suggesting that bile acids induced cell death was not necroptosis (**Figures S1N and S1O**). Moreover, our previous results excluded TLR4 dependence of bile acids function(Hao et al., 2017) and the assay here further verified that bile acids applied as well as cell cultures were free of LPS contamination and bile acids were tested without LPS priming for all experiments performed in this study (**STAR Methods**).

### GSDME Is Involved in Bile Acids Induced Pyroptosis

There are two key executors known to trigger pyroptosis, including GSDMD and GSDME. GSDMD is necessary for caspase-1/4/8/11-induced pyroptosis(Feltham and Vince, 2018; Lee et al., 2018; Shi et al., 2015) and caspase-3 was recently shown to cleave GSDME (Rogers et al., 2017; Wang et al., 2017). Unexpectedly, bile acids activated caspase-11 but were incapable of cleaving GSDMD to a p30 fragment which is essential for pore formation in LPS-induced cell lysis(Shi et al., 2014). Instead, GSDMD was found cleaved to a p43 fragment that was performed by caspase-3 in bile acids treated cells (**Figures 1F**). Of interest, bile acids triggered the cleavage of GSDME to a p30 fragment and is dependent on caspase-11 (**Figures 1F and 1M**). BMDMs isolated from *GSDME* rather than *GSDMD* knockout mice were resistant to bile acids induced cell death, LDH release and IL-1α secretion (**Figures 2A-D**). Moreover, *GSDME*^-/-^ BMDMs were resistant to bile acids induced pyroptosis (**Figures 2E-F**). Bile acids also induced a GSDME dependent pyroptosis in HepG2 cells since *GSDME* silenced cells were insensitive to bile acids induced pyroptosis (**Figure S2**). Bile acids induced caspase-3 activation and cleavage of GSDME were found absent in *caspase-11^-/-^* BMDMs (**Figure 1M**, **Figures 2G and 2H**). Moreover, in the *in vitro* cell free system, activated caspase-4 directly cleaved and activated caspase-3 (**Figures 2I and 2J**). These results suggest that caspase-11/4 acts as upstream of caspase-3 that cleaves GSDME leading to pyroptosis upon bile acid stimulation. Notably, LPS promoted pro-caspase-4 interacting with GSDMD, and in the cell free system of LPS/caspase-4, caspase-3 was unable to compete with GSDMD to be activated by caspase-4 (**Figure 2K**), supporting that GSDMD is recruited to LPS/caspase-4 complex before its cleavage by caspase-4. In contrast, pro-caspase-4, in conditions of bile acids activation, could not bind with GSDMD (**Figures 2L**), and GSDMD was instead erroneously cleaved by caspase-3 to produce a p43 fragment (**Figures 2M**). These results indicate that the direct binding of pro-caspase-11/4 with GSDMD upon LPS ligation is a prerequisite for the subsequent caspase-11/4 oligomerization and activation and thereby correctly cleaving GSDMD, while caspase-4 activated by bile acids instead cleaves caspase-3 to trigger a GSDME-mediated pyroptosis.

**Figure 2.**
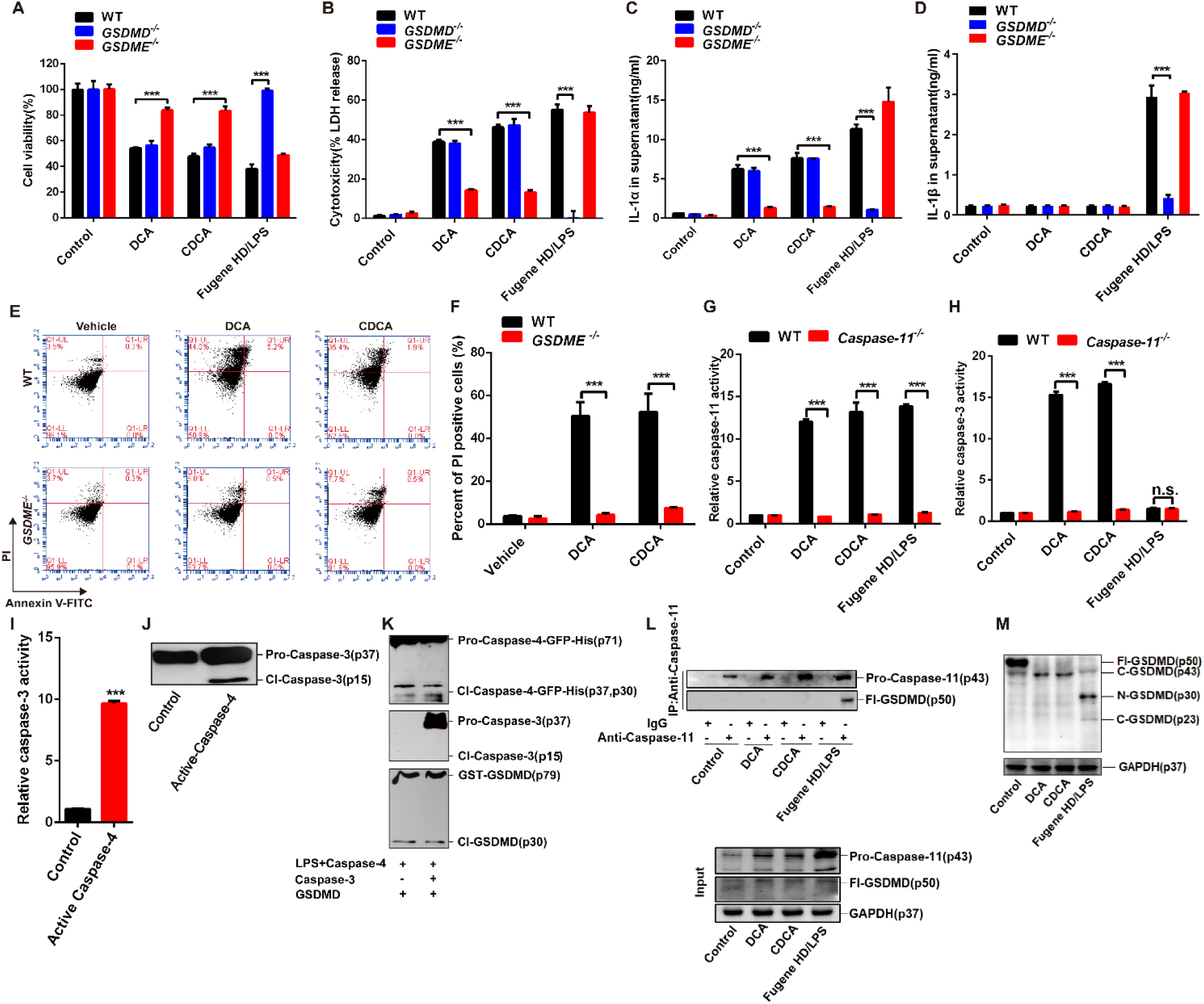
Bile Acids Activate A Caspase-4/11-Caspase-3-GSDME Pyroptotic Pathway. **(A-D)** CCK8 assay for cell viability **(A)**, LDH release for cytotoxicity **(B)** and IL-1α/β secretion **(C and D)** of BMDMs isolated from WT, *GSDMD^-/-^* and *GSDME^-/-^* mice. Experimental design as in Figure 1A-D. **(E)** Flow cytometry analysis of Annexin V-FITC/PI staining in WT and *GSDME^-/-^* BMDMs treated with 200 μM DCA or CDCA for 4 hrs. **(F)** Proportions of PI positive cells, quantified from images as shown in (E). **(G and H)** Caspase activities in WT and *caspase-11^-/-^* BMDMs were analyzed by using Caspase Colorimetric Assay Kit (Biovision). Cells were treated with 200 μM bile acid for 1hr; for LPS group, LPS-primed cells were transfected with 5 μg/ml LPS by 0.3% (v/v) Fugene HD for 9 hrs. **(I and J)** 200 μM recombinant human pro-caspase-3 was incubated with 2 U (relevant to ∼ 100 nM) recombinant active caspase-4 for 2 hrs at 37 °C in a 50 μL reaction system. Then caspase-3 activation was analyzed by Ac-DEVD-ρNA **(I)** and immunoblotting **(J)** **(K)** 1 mg/mL LPS was incubated with 1 μM recombinant human pro-caspase-4, −3 and GSDMD in a reaction system for immunoblot analysis of active-caspase-3 and GSDMD fragments. **(L)** Co-immunoprecipitation analysis of pro-caspase-11 and full length GSDMD (Fl-GSDMD) in BMDMs lysates. Experimental design as in Figure 1A-D. **(M)** Immunoblot analysis of p43, p30, p23 fragments GSDMD in cell lysates of BMDMs at high exposure. Experimental design as in Figure 1A-D. GAPDH was used as loading control in immunoblot analyses of cell lysates. Bar graphs expressed as mean ± SEM (n=3). \*\*\**p*<0.001; n.s., non-significant difference compared to control unless indicated otherwise. See also **Figure S2**.

### MPT Determines Bile Acids Induced Pyroptosis

A critical question is how caspase-4 detects bile acids signal to trigger pyroptosis. LPS transfected to the cytoplasm directly binds with caspase-4/11 inducing oligomerization and activation for triggering pyroptosis(Shi et al., 2014). We asked whether bile acids can also directly bind and activate caspase-4. Microscale thermophoresis (MST) analysis showed that bile acids, unlike LPS, could not directly bind with recombinant human caspase-4 nor murine caspase-11 (**Figure S3A**). In cell free system, LPS, but not bile acids, directly activated caspase-4 and −11 (**Figures S3B and S3C**). These results indicate that bile acids activate caspase-4 in a manner distinct to that of LPS, hinting for that certain cell-intrinsic factors activated by bile acids might be involved in activation of caspase-4.

Mitochondrion is a pivotal organelle in controlling cell fates and bile acids are known for their effects in inducing mitochondrial stress. We hypothesized that mitochondrion might be involved in regulation of pyroptosis. Bile acids treatment at concentrations capable of triggering pyroptosis markedly induced MPT as evidenced from the loss of mitochondrial cristae and calcein AM/CoCl2 assay in BMDMs (**Figures 3A-C**). MPT is a state characterized with abrupt loss of integrity of inner mitochondrial membrane and thus allowing free permeability to small solutes resulted in the loss of osmotic balance(Baines et al., 2005; Nakagawa et al., 2005). Time-course assay indicated bile acids induced a long-lasting MPT as characterized by persistent dissipation of mitochondrial transmembrane potential (ψ and loss of integrity of inner mitochondrial membrane (**Figures 3D and 3E**). Persistent MPT is mediated by the irreversible opening of a multiprotein complex pore, called MPTP (Izzo et al., 2016). Although the exact molecular components of MPTP remain unclear and may vary in a context dependent manner, several proteins including adenine nucleotide translocator (ANT), cyclophilin D (CYPD) and voltage-dependent anion channel (VDAC) have been considered as functional components(Baines et al., 2005; Javadov et al., 2017; Karch and Molkentin, 2014; Nakagawa et al., 2005; Zhivotovsky et al., 2009). We screened a panel of inhibitors to these MPTP components. Among of these, bongkrekic acid (BKA), a selective ANT inhibitor, significantly protected cells from bile acid-induced MPT, cell death, and LDH release, and abolished the activation of caspase-11/4, caspase-3 and GSDME (**Figures 3F-M, Figures S3D and S3E**). In contrast, BKA could not inhibit LPS induced LDH release, cell death, and secretion of IL-1α and IL-1β (**Figures S3F-I**). ANT is responsible for translocating ATP/ADP across the inner mitochondrial membrane (Clemencon et al., 2013; Halestrap and Brenner, 2003). Notably, bile acids induced MPT was featured with an immediate release of ATP to the cytoplasm within 30 min post bile acids treatment (**Figure 3N**). Cytoplasmic ATP levels decreased to below the basal level (**Figure 3N**) from 60 min post bile acids treatment, in line with previous reports that MPT inhibited ATP generation(Cogliati et al., 2013). All these results support that bile acids induced pyroptosis may be initiated by ANT-facilitated MPT.

**Figure 3.**
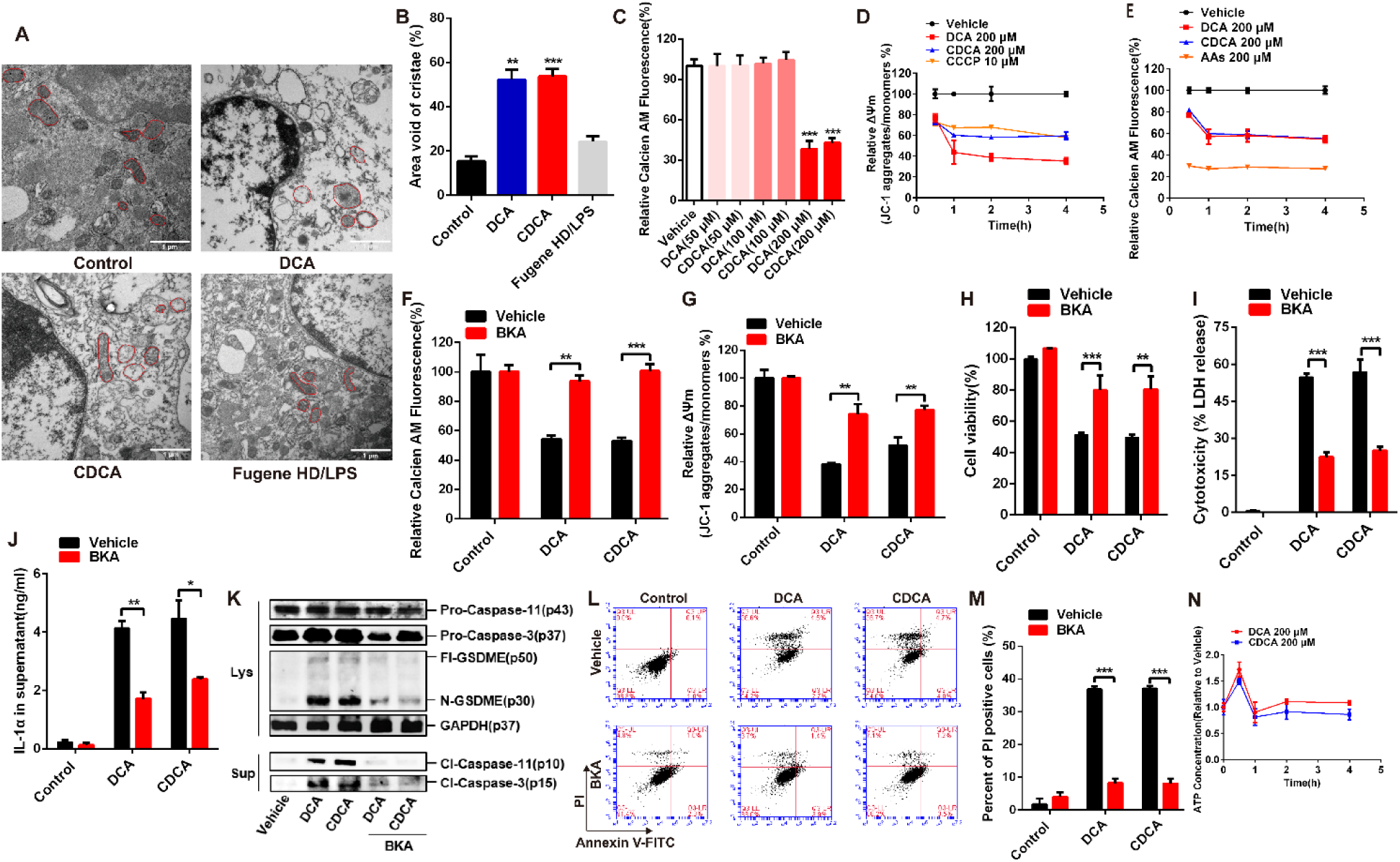
Bile Acids Triggered Pyroptosis Is Dependent of Permanent MPT. **(A)** Representative electron microscopy images of BMDMs; representative mitochondrial morphologies were marked with red lines. Scale bars, 1 μm. **(B)** Mitochondrial cristae density was represented as percent area void of cristae (minimum of 40 mitochondria were analyzed per well; triplicate wells per group); quantified from images as shown in (A). **(C)** MPT was detected by calcien-AM/CoCl_2_ assay; BMDMs were treated with 50-200 μM bile acid for 1 hr; data are relative to vehicle group. **(D and E)** BMDMs were treated with 200 μM bile acid; time-course analysis of MPT were detected by calcien-AM/CoCl_2_ assay and dissipation of Δψ_m_ by JC-1 assay; carbonyl cyanide 3-chlorophenylhydrazone (CCCP) and arachidonic acids (AAs) were used as positive controls of Δψ_m_ dissipation and MPT respectively. **(F)** MPT was detected by calcien-AM/CoCl_2_ assay; BMDMs were pre-treated with or without 20 μM ANT inhibitor BKA for 2 hrs, then treated with 200 μM bile acids. **(G)** Δψ_m_ dissipation indicated by JC-1 disaggregation, BMDMs were stimulated with 200 μM BAs for 1hr. **(H-J)** CCK8 assay for cell viability **(H)**; LDH release for cytotoxicity **(I)** and IL-1α release **(J)** in BMDMs; cells were pretreated with or without 20 μM BKA for 2 hrs followed by 200 μM DCA or CDCA for 4 hrs. **(K)** Immunoblots of caspase-11, caspase-3, GSDME and GAPDH in BMDMs. Experimental design as in (H-J). **(L)** Flow cytometry analysis of Annexin V/PI stained BMDMs. Experimental design as in (H-J). **(M)** Proportions of PI positive cells; quantified from images as shown in (L). **(N)** Fold change of cytoplasmic ATP concentration in BMDMs treated with 200 μM bile acid at indicated times. GAPDH was used as loading control in immunoblot analyses of cell lysates. Bar graphs expressed as mean ±SEM (n=3). *, *p*<0.05; **, *p*<0.01; ***, *p*<0.001; n.s., non-significant. See also **Figure S3**.

### Caspase-4/11 Is A General Sensor of MPT

Previously, MPT induced cell death was defined as regulated necrosis but with unclear molecular events(Baines et al., 2005; Ying and Padanilam, 2016). Our results indicated that MPT triggered by bile acids resulted in caspase-4 and GSDME mediated pyroptosis. We next asked whether caspase-4 represents a general sensor of MPT and whether MPT-elicited regulated necrosis was typically caspase-4 mediated pyroptosis. As expected, mitochondrial calcium overload induced by thapsigargin (TG) provoked typical MPT and triggered pyroptotic cell death which was blocked by BKA (**Figures 4A and 4B, Figures S4A-E**). Moreover, TG induced pyroptotic cell death and release of LDH and IL-1α were mediated by caspase-11, since BMDMs isolated from *Caspase-11*^-/-^ mice were insensitive to TG treatment (**Figures 4C-E**). More specifically, lonidamine (LND), an adjuvant anticancer agent targeting ANT1(Belzacq et al., 2001), induced a persistent MPT, and consequently pyroptotic cell death characterized with activation of caspase-11 and caspase-3, GSDME cleavage, and increased release of IL-1α and LDH in BMDMs in a ANT1 dependent manner (**Figures 4F-K, Figures S4F-J**). LND and bile acid also induced pyroptotic cell death in HepG2 cells, which was largely abolished by BKA pretreatment and silencing *ANT1* (**Figures 4L-R**). These results support that caspase-4 and GSDME mediated pyroptosis represents a general mechanism underlying MPT induced regulated necrotic cell death.

**Figure 4.**
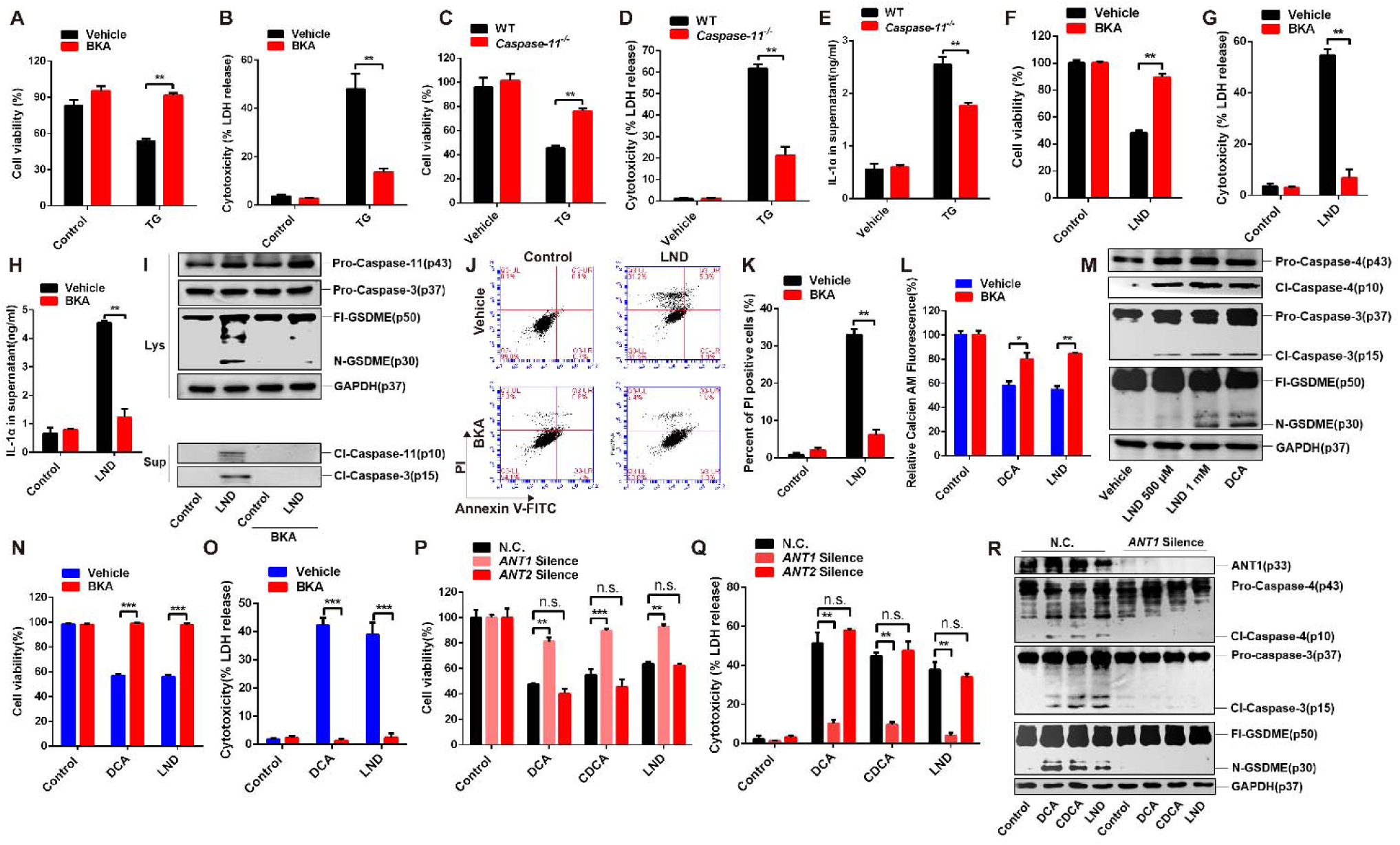
Caspase-4/11 Elicited Pyroptosis Dominants in MPT-Triggered Cell Death. **(A and B)** CCK8 assay for cell viability **(A)**; LDH release for cytotoxicity **(B)** of BMDMs; cells were pre-treated with 20 μM BKA for 2 hrs before 20 μM TG challenge for 1 hr. **(C-E)** CCK8 assay for cell viability **(C)**; LDH release for cytotoxicity **(D)** and IL-1α release **(E)** in BMDMs from WT and *caspase-11*^-/-^ mice; cells were treated with 20 μM TG for 1 hr. **(F-H)** CCK8 assay for cell viability **(C)**; LDH release for cytotoxicity **(D)** and IL-1α release **(E)** in BMDMs; cells were pre-treated with 20 μM BKA for 2 hrs followed by 1 mM LND for 12 hrs. **(I)** Representative immunoblots of caspase-11, caspase-3, GSDME and GAPDH in cell lysates (Lys) and supernatant (Sup) of BMDMs; experimental design as in (F-H). **(J)** Flow cytometry analysis of Annexin V/PI stained BMDMs; experimental design as in (F-H). **(K)** Proportions of PI positive cells; quantified from images as shown in (J). **(L)** MPT was detected by calcien-AM/CoCl_2_ assay; HepG2 cells were pre-treated with 20 μM BKA for 2hrs followed by 1 mM LND for 4 hrs or 200 μM DCA for 1 hr. **(M)** Representative immunoblots of caspase-4, caspase-3, GSDME and GAPDH in cell lysates of HepG2 cells; cells were treated with 500μM/1 mM LND for 12 hrs or 200 μM DCA for 4 hrs. **(N and O)** CCK8 assay for cell viability **(N)** and LDH release for cytotoxicity **(O)** in HepG2 cells; cells were pre-treated with 20 μM BKA for 2hrs followed by 1 mM LND for 12 hrs or 200 μM DCA for 4hrs. **(P and Q)** CCK8 assay for cell viability **(P)** and LDH release for cytotoxicity **(Q)** in *ANT1/ANT2* silenced HepG2 cells compared with negative control (N.C.); cells were treated with 200 μM DCA/CDCA for 4 hrs or 1 mM LND for 12 hrs. **(R)** Representative immunoblots of ANT1, caspase-4, caspase-3, GSDME and GAPDH in cell lysates of *ANT1* silenced HepG2 cells compared with N.C.; experimental design as in (P and Q). GAPDH was used as loading control in immunoblot analyses of cell lysates. Bar graphs expressed as mean ±SEM (n=3). *, *p*<0.05; **, *p*<0.01; ***, *p*<0.001; n.s., non-significant. See also **Figure S4**.

### Apaf-1/Caspase-4/11 Pyroptosome Determines MPT Triggered Pyroptosis

MPT activates a caspase-4/caspase-3/GSDME pyroptotic pathway, which is distinct to both LPS induced caspase-4/GSDMD and chemotherapeutic drugs triggered caspase-3/GSDME pathway. Classical chemotherapeutic drugs were incapable of inducing MPT and caspase-4 activation (**Figures S5A-D**). We were thus of particular interest of the mechanism underlying intrinsic activation of caspase-4 by MPT. It has been recognized that oligomerization of precursor molecules is a general mechanism for the activation of both apoptotic and inflammatory caspases. Caspase-9 and caspase-1 is activated on a protein complex platform known as apoptosome and canonical inflammasome, respectively. We hypothesized that the activation of caspase-4/11 may also necessitates a protein complex as a scaffold. To this end, we performed an IP assay using caspase-11 antibody to determine possible interacting proteins. Surprisingly, Apaf-1, the key adapter in intrinsic apoptosis pathway(Hu et al., 2014; Hu et al., 1998; Pang et al., 2015), was co-precipitated with caspase-11 in BMDMs (**Figure 5A**). Caspase-9 was slightly co-precipitated with caspase-4 observed from high exposure immunoblots, possibly due to cross reaction between caspase-9 antibody with caspase-4. Direct specific interaction of Apaf-1 with caspase-4 was confirmed by His-Tag pull down assay in HEK293T cell line and MST assay (**Figures 5B-D**), while caspase-9 was not witnessed from this His-Tag pulldown assay using caspase-4 antibody, supporting that caspase-9 is unlikely involved in MPT-triggered assembly of protein complex. The assembly of Apaf-1-caspase-4 protein complex was further visualized by confocal analysis in HepG2 cells treated with bile acids and LND (**Figure S5E**).

**Figure 5.**
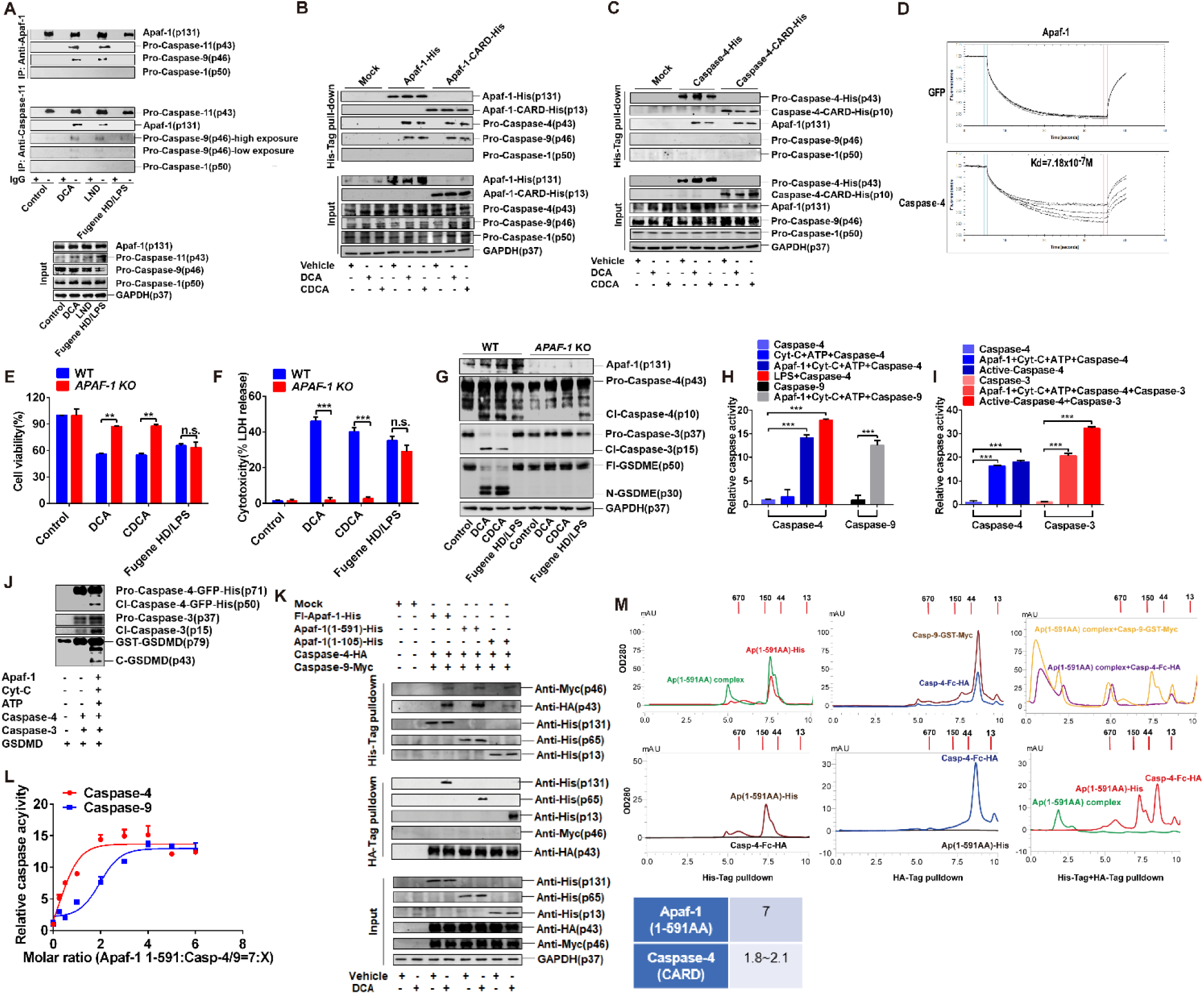
MPT But not MOMP Promotes Apaf-1 Pyroptosome Assembly. **(A)** Co-immunoprecipitation analysis of Apaf-1, caspase-11, −9, −1 from lysates of BMDMs; cells were treated with 200 μM DCA for 30 min, 1 mM LND for 6 hrs, or for LPS group, LPS-primed BMDMs were transfected with 5 μg/mL LPS by 0.3% (v/v) Fugene HD for 6 hrs. **(B)** His-Tag pull-down analysis of the interaction between Apaf-1-His or Apaf-1-CARD-His with endogenous pro-caspase-4, −9, −1. His tagged full length Apaf-1 or Apaf-1-CARD overexpressing plasmid were transfected into HepG2 cells, then cells were treated with 200 μM DCA or CDCA for 30 min. **(C)** His-Tag pull-down analysis of the interaction between pro-caspase-4-His or caspase-4-CARD-His with endogenous Apaf-1, pro-caspase-9, −1. His tagged full length caspase-4 or caspase-4-CARD overexpressing plasmid were transfected into HepG2 cells, experimental design as in (B). **(D)** MST measurements of direct binding between Apaf-1 and caspase-4. Apaf-1 concentrations at 23, 46, 92, 184, 370, or 740 nM (from bottom to top) were pre-incubated with cytochrome-c (Cyt-C, half of Apaf-1 concentrations) and 5 mM ATP overnight at 4 °C to achieve active Apaf-1 complex. 1 μM recombinant pro-caspase-4-eGFP was pre-labeled and incubated with Apaf-1 complex for 30 min at room temperature, then inhaled with capillaries for MST assay. eGFP was included as negative control. **(E and F)** CCK8 assay for cell viability **(E)** and LDH release for cytotoxicity **(F)** in *APAF-1* knockout HepG2 cells compared with WT group. Experimental design as in Figure 1A-D. **(G)** Representative immunoblots of Apaf-1, caspase-4, −3, GSDME and GAPDH in lysates of *APAF-1* knockout HepG2 cells compared with WT group. Experimental design as in Figure 1A-D. **(H and I)** Caspase activities in cell-free system were determined by substrate Ac-LEVD-ρNA for caspase-4, Ac-DEVD-ρNA for caspase-3 and Ac-LEHD-pNA for caspase-9, respectively. 400 nM Apaf-1 complex were pre-constructed as in (D), then pro-caspases were added at 1 μM; 2 U active-caspase-4 incubated with caspase-3 and 1 mg/ml LPS incubated with caspase-4 were included as positive controls; all groups were incubated at 37 °C for 2 hrs. **(J)** Immunoblotting analysis of caspase-4, −3 and GSDMD activation in recombinant protein incubation system. 400 nM Apaf-1 were pre-incutabed with 200 nM Cyt-C and 5 mM ATP overnight, then 1 μM recombinant pro-caspase-4 were added and incubated at 37 °C for 2 hrs, followed by 200 nM caspase-3 and 200 nM GSDMD for further 1hr at 37 °C. **(K)** Pull-down analysis of interaction of caspase-4-HA/caspase-9-Myc with full length (Fl)-Apaf-1-His, Apaf-1(1-591AA)-His and Apaf-1(1-105)-His in HEK293T cells. Cells were transfected with indicated plasmids, then treated with 200 μM DCA for 30 min. **(L)** Relative caspase activities in constructed protein system were detected by substrate Ac-LEVD-pNA for caspase-4 and Ac-LEHD-pNA for caspase-9. 0.7 μM of Apaf-1(1-591AA) was incubated with 5 mM ATP and gradient concentrations of caspase-4 or −9 (0.025, 0.05, 0.1, 0.2, 0.3, 0.4, 0.5, 0.6 μM) for 2 hrs at 37 °C. The relative caspases activities were measured against the molar ratios of Apaf-1 (1-591AA) to Caspase-4/9 (X represents Apaf-1 : Caspase=7:X). **(M)** Size exclusion chromatograms analysis of Apaf-1(1-591AA)/caspase-4-CARD complex composition. Stable complex of Apaf-1(1-591AA)-His and caspase-4-CARD-Fc-HA were constructed (4 μM : 2 μM) as in (L), then purified by double pull-down of His-tag and HA-Tag; native complex or denatured components were subjected to SEC analysis and peak area of Apaf-1 (1-591AA)-His and Casp-4-Fc-HA at OD280nm were used for ratio calculation. GAPDH was used as loading control in immunoblot analyses of cell lysates. Bar graphs expressed as mean ±SEM (n=3). **, *p*<0.01; ***, *p*<0.001; n.s., non-significant. See also **Figure S5**.

Next, we asked whether Apaf-1 is functionally involved in MPT triggered pyroptosis. Bile acids and LND were unable to activate caspase-4/caspase-3/GSDME signaling, pyroptotic cell death, and LDH release in *APAF-1* knockout HepG2 cells(**Figures 5E-G Figures S5F-H**). Moreover, the Apaf-1/caspsae-4 protein complex assembly was blocked by BKA pretreatment (**Figure S5I**). These results indicate that Apaf-1 is essential for caspase-4 recruitment and activation. To verify the Apaf-1-caspase-4 complex is both sufficient and necessary for activation of pyroptotic signals, we reconstituted a molecular system containing recombinant human Apaf-1 and caspase-4, in which cytochrome C and ATP were included to activate the adapter Apaf-1. In contrast to LPS that directly bound to and activated caspase-4, Apaf-1 activated caspase-4 only in the presence of ATP and cytochrome C as cofactors (**Figures 5H and 5I**), which is quite similar to the mode of caspase-9 activation on the apoptosome scaffold. Pro-caspase-3 added into this reconstituted Apaf-1-caspase-4 system could be activated (**Figures 5I and 5J**), supporting that this system is functionally active and that caspase-4 acts upstream of caspase-3. Therefore, we termed MPT triggered assembly of such a protein complex as Apaf-1 pyroptosome to which caspase-4 was recruited and activated to facilitate pyroptosis.

Caspase-9 is recruited to Apaf-1 via multimeric caspase recruitment domain (CARD): CARD interactions. It is thus reasonable to speculate that caspase-4 may also interact with Apaf-1 via CARD:CARD interactions. His-tagged recombinant human Apaf-1 plasmids (including full-length, 1-591AA representing a functional Apaf-1 that could self-oligomerization without cytochrome C, and 1-105AA representing the CARD domain) were transfected into HEK293T cells. Upon bile acid treatment, all three Apaf-1 fragments showed marked interaction with caspase-4(**Figure 5K**). To provide direct evidence, a fixed amount of preassembled Apaf-1 (1-591AA) was incubated with different concentrations of caspase-4 or caspase-9 (Apaf-1: caspase molar ratios range from 7:0.25 to 7:6), and then the protease activity of caspase-4/9 was measured and plotted against the molar ratios of caspase-4/9 over Apaf-1 (1-591AA). Caspase-4 activity reached the maximum at the molar ratio of 2:7 in contrast with 4:7 for caspase-9 (**Figure 5L**). Moreover, Kcat/Km of caspase-4 was ∼22% higher than that of caspase-9 (**Figures S5J and S5K**). To further determine the stoichiometric ratio of Apaf-1 and caspase-4 in pyroptosome, Apaf-1(1-591AA)-His and caspase-4(CARD)-Fc-HA were incubated to form a stable complex. This complex was then isolated via double purification by HA-tag and His-Tag pulldown and quantified by size exclusion chromatography. The molar ratio of Apaf-1(1-591AA) to caspase-4 (CARD) was calculated to be 3.4∼3.9 in three independent experiments, which corresponds to 1.8∼2.1 molecules of caspase-4 (CARD) for each Apaf (1-591) heptamer (**Figure 5M**). Together, these results support a 7:2 molar ratio between Apaf-1 and caspase-4 in the pyroptosome platform, which is different from that of the apoptosome stoichiometry of 7:4 between Apaf-1 and caspase-9 (**Figure 5L**)(Hu et al., 2014).

### Mitochondrion Quantitatively Dictates Apoptosis and Pyroptosis

Notably, caspase-4 was activated by Apaf-1 in a similar mode to that of caspase-9. A critical question is how cells respond differentially to mitochondrial stress and thereby dictating the choice between apoptosis and pyroptosis. Bile acids at 50 μM were sufficient for inducing MOMP-triggered Apaf-1/caspase-9 assembly and thereby activating caspase-9 and caspase-3. By contrast, MPT-triggered activation of caspase-11/caspase-3 was achieved only at high dose up to 200μ **(Figures 6A-G)**, suggesting a higher threshold is necessary for caspase-11/4 than that for caspase-9 recruitment and activation. Moreover, the time-course assay clearly showed that MOMP induced by low dose bile acids and doxorubicin (DOX) activated caspase-9 and −3 from 4 hrs post-treatment, while MPT induced by high dose bile acids triggered an immediate activation of caspase-11 peaking at 1 hr after treatment. Notably, caspase-9 was not activated from the high dose bile acids treatment. HepG2 cells treated with 200 μM bile acid also underwent a caspase-4 mediated pyroptotic cell death. Time-course assay of enzyme activities indicated that the activation of caspase-4 and −3 induced by bile acids treatment was much stronger than that of caspase-9 and −1 (**Figures S6A and S6B**). Knockout of *CASPASE-4*, but not *CASPASE-9*, significantly reduced pyroptotic cell death as evidenced by cell viability, LDH release, IL-1α secretion and PI positive staining cells (**Figure S6C-I**). Because both apoptosis and pyroptosis depend on mitochondrial release of cytochrome C for the activation of Apaf-1, we asked how this molecular event is differently regulated between apoptosis and pyroptosis. To this end, a time-course analysis of cytochrome C and ATP release were performed. MPT led to an earlier increase of cytoplasmatic cytochrome C in comparison with that by MOMP (**Figure 6H**). MPT, but not MOMP, induced an immediate release of ATP to the cytoplasm (**Figure 6I**). We supposed the energy threshold for caspase-11/-4 recruited to Apaf-1 might be much higher than that for apoptosome assembling. We further tested the ATP threshold in a cell free incubation system using recombinant human Apaf-1(1-591AA). Caspase-4 recruited to Apaf-1 necessitated a much higher ATP level than that of caspase-9. Of interest, caspase-4 competitively inhibited caspase-9 activation at conditions with high ATP level (≥2mM) (**Figure 6J**), which was in line with the time-course assay of enzyme activities (**Figures 5C and 5D**). Together, these results indicate that the releasing rate and amount of cytochrome C and ATP may represent a key threshold dictating the choice between apoptosis and pyroptosis. An abrupt release of large amount of cytochrome C and ATP in conditions of MPT triggered by sever mitostess favors pyroptosome assembly and caspase-4/11 activation for facilitated GSDME-executed pyroptosis. In contrast, MOMP resulted from mild mitostress preferentially promotes Apaf-1 apoptosome assembly for initiation of caspase-9 mediated apoptosis (**Figure 6K**). Moreover, MPT triggered caspase-4 activation initiated pyroptosis is much quicker and fulfilled in a much shorter time scale than that of MOMP prompted caspase-9 activation-initiated apoptosis.

**Figure 6.**
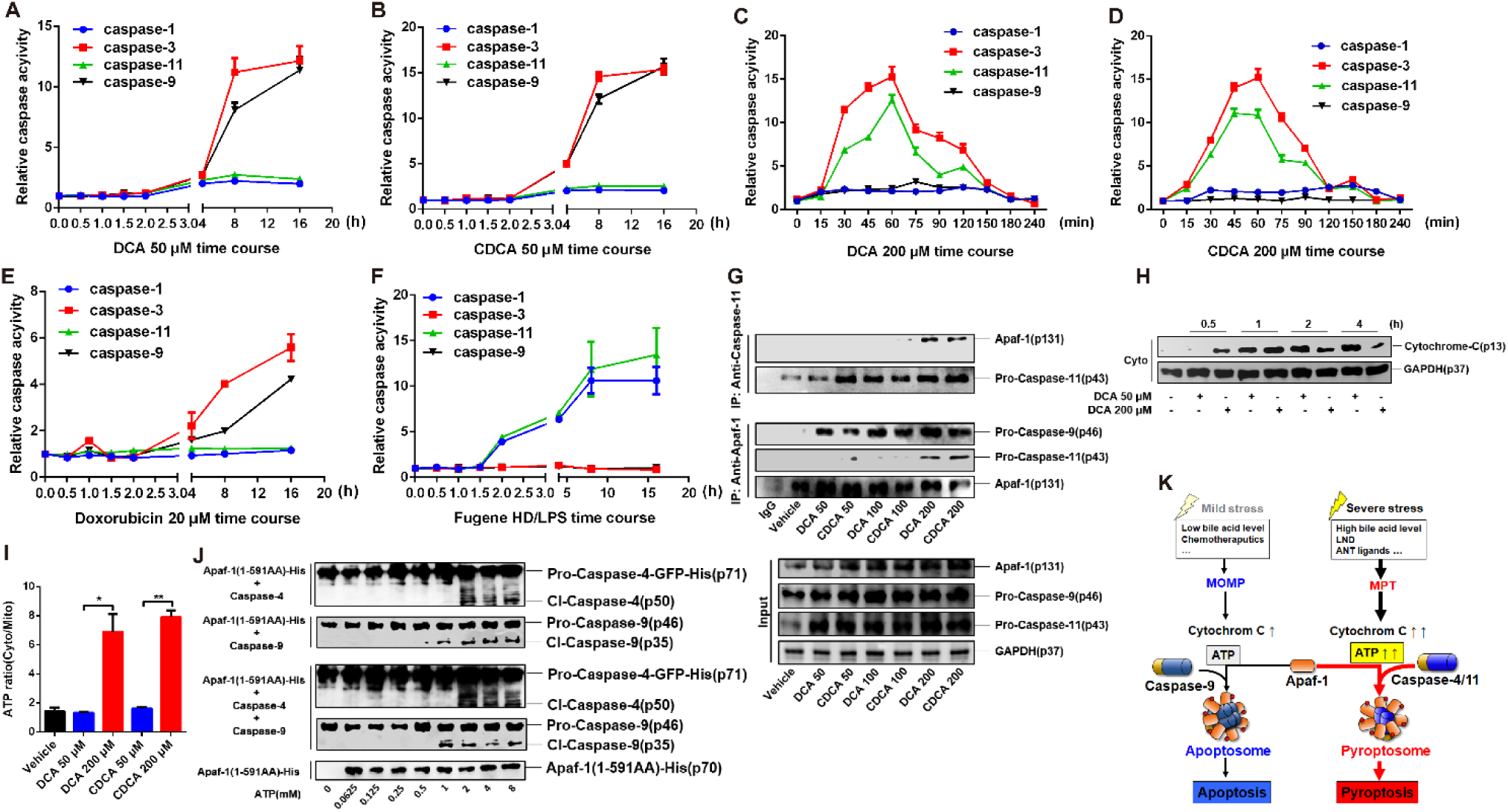
Mitochondria Quantitatively Dictate Apaf-1 Apoptosome and Pyroptosome Assembly. **(A-F)** Time course analysis of fold change of caspase activities in lysates of BMDMs were measured by use Caspase Activity Assay Kits of caspase-1, −3, −11, −9 respectively. Cells were treated with 50 or 200 μM bile acids, 20 μM Doxorubicin, or for LPS group, LPS-primed cells were transfected with 5 μg/mL LPS by Fugene HD for indicated hrs. **(G)** Co-immunoprecipitation analysis of interaction between Apaf-1 and pro-caspase-9, −11 in lysates of BMDMs; cells were treated with 50∼200 μM bile acids for 30 min. **(H)** Representative immunoblots of Cytochrome-C and GAPDH in cytoplasmic lysates (Cyto) of BMDMs treated with 50 or 200 μM DCA for indicated hrs,. **(I)** Relative level of ratio of cytoplasmic ATP concentration to mitochondrial ATP concentration in BMDMs; cells were treated with 50 or 200 μM DCA for 30 min. **(J)** Activation of caspase-4 and −9 in cell free systems were indicated by immunoblotting. 400 nM Apaf-1(1-591AA) was incubated with 0∼8 mM ATP at 4 °C overnight, then 200 nM caspase-4 and/or −9 were added and further incubated at 37 °C for 2 hrs. **(K)** Illustration of dictations between pyroptosome and apoptosome assembling. GAPDH was used as loading control in immunoblot analyses of cell lysates. Bar graphs expressed as mean ±SEM (n=3). *, *p*<0.05; **, *p*<0.01. See also **Figure S6**.

### Caspase-11/GSDME Mediated Pyroptosis Dictates Cholesteric Liver Injury

Our study identified a cell-intrinsic signal pathway that activates caspase-4 to elicit pyroptosis under sterile conditions. We thus asked whether such a cell-intrinsic pyroptosis triggered by MPT is pathophysiologically relevant. Since bile acids at pathologically high levels activate MPT leading to pyroptosis, we employed a typical cholestasis model by bile duct ligation (BDL). BDL mice were characterized with acute liver injury and drastic accumulation of bile acids in both circulating system and the liver. Here we found BDL mice were characterized with significant upregulation of ANT1 and persistent MPT as assessed by loss of mitochondria cristae (**Figures 7A and 7B, Figure S7A**). The increase of serum IL-1α β HMGB1 levels was observed at an early stage of 7 days after BDL (**Figures S7B-D**). The formation of Apaf-1/caspase-11 pyroptosome complex, caspase-11 activation and the cleavage of caspase-3 and GSDME were witnessed **(Figures 7C and 7D)**. Notably, caspase-1 and downstream IL-1β were also activated in BDL mice (**Figure 7E, Figure S7C**), which is in line with our previous results(Hao et al., 2017). Consistently, GSDMD was partially cleaved to the pore-forming p30 fragment by caspase-1 concurrently with inactive p43 fragment by capase-3 (**Figure 7E**). *Caspase-11^-/-^* mice were resistant to BDL induced activation of caspase-3, GSDME cleavage, and secretion of IL-1α/β and HMGB1 (**Figures 7F-I**). Moreover, BDL induced mortality was completely rescued in *GSDME^-/-^* mice (**Figures 7J-M**). Although the contribution of caspase-1/GSDMD signal cannot be excluded, these results indicate that caspase-11-caspase-3-GSDME induced pyroptotic cell death is dominant in determining BDL induced cholesteric liver failure.

**Figure 7.**
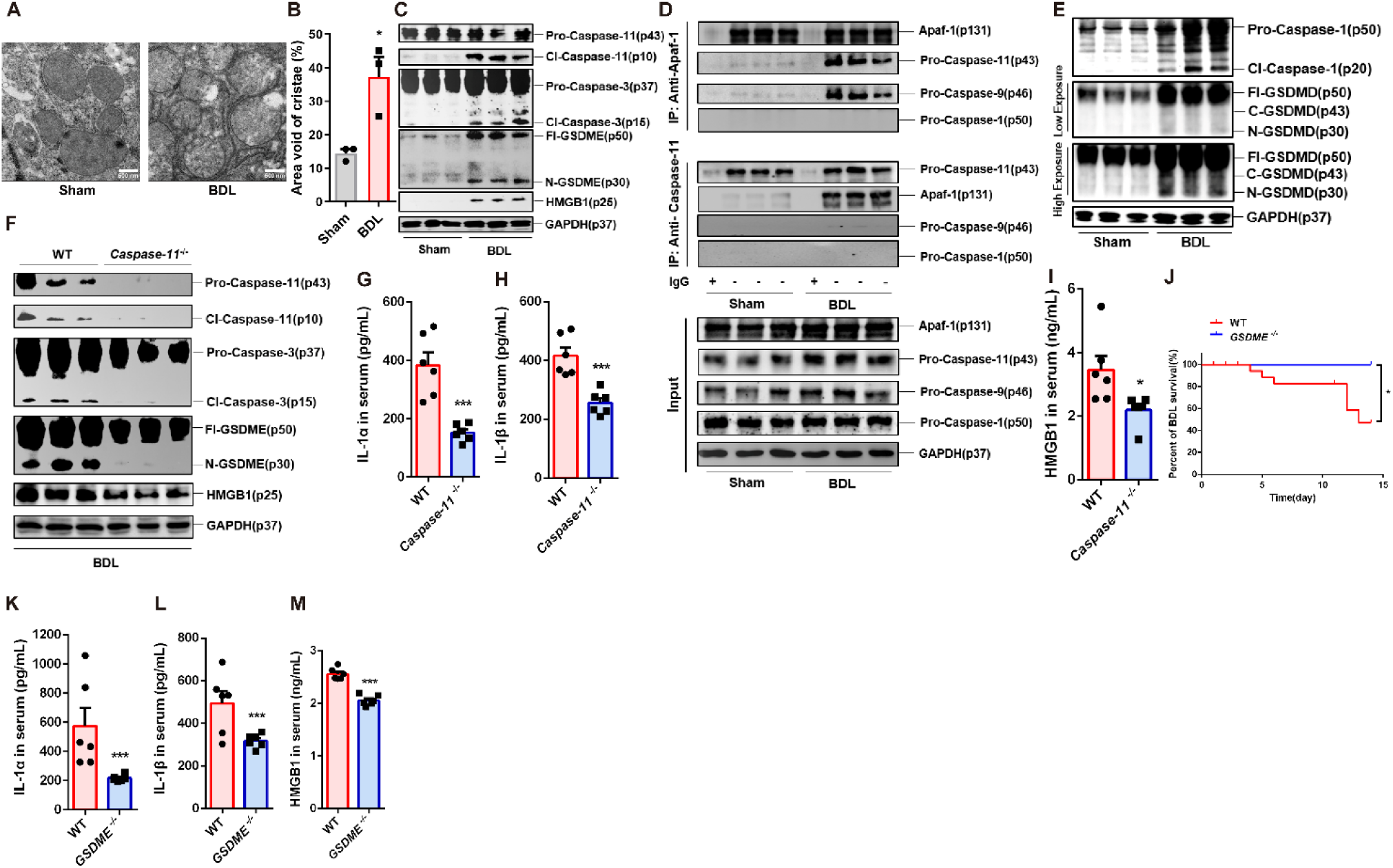
Caspase-11 and GSDME Dependent Pyroptosis Dominants BDL Induced Cholesteric Liver Failure. **(A)** Representative electron microscopy images of mitochondrial swelling from liver of mice 7 days after BDL or sham (n=3), scale bars, 500 nm. **(B)** Mitochondrial cristae loss and decrement of electron density, represented as percent area void of cristae, quantified from images as shown in (A); minimum of 40 mitochondria were analyzed per mouse (n=3). **(C)** Representative immunoblots of caspase-3, −11, GSDME, HMGB1 and GAPDH in liver lysates of mice 7 days after BDL/sham (n=3). **(D)** Co-immunoprecipitation analysis of interaction between Apaf-1 and caspase-1/9/11 in liver lysates of mice 7 days after BDL/sham (n=3). **(E)** Representative immunoblots of caspase-1, GSDMD and GAPDH in liver lysates of mice 7 days after BDL/sham (n=3). **(F)** Representative immunoblots of caspase-3, −11, GSDME, HMGB1 and GAPDH in liver lysates of WT and *caspase-11^-/-^* mice 7 days after BDL/sham (n=3). **(G-I)** Serum levels of IL-1α, IL-1β and HMGB1 in WT and *caspase-11*^-/-^ mice 7 days after BDL/sham (n=6). **(J)** Survival analysis of WT and *GSDME*^-/-^ mice after BDL (n = 10). **(K-M)** Serum levels of IL-1α, IL-1β and HMGB1 in WT and *GSDME*^-/-^ mice 14 days after BDL/sham (n=6). GAPDH was used as loading control in immunoblot analyses of tissue lysates, lysates from six mice was merged by two-into-one to n=3. Scatter plot with box graphs expressed as mean ± SEM. *, *p*<0.05; ***, *p*<0.001. See also **Figure S7**.

## DISCUSSION

Caspase-4/11 is a sensor of intracellular LPS, explaining why microbial infections trigger pyroptotic cell death. However, it remains elusive whether and how caspase-4/11 detect endogenous derived factors for eliciting pyroptosis implicated in diverse sterile diseases. Here, we establish a causal link of caspase-4/11 activation and pyroptosis under sterile conditions. Upon MPT stimulation, cytochrome c and ATP released into the cytoplasm promote the assembly of pyroptosome consisted of Apaf-1 and caspase-4/11 and thereafter the activation of caspase-3 that cleaves GSDME for triggering pyroptosis.

MPT can be induced by calcium overload, oxidative stress, and the accumulation of some toxic endobiotics like bile acids, the conditions of which are implicated in many kinds of diseases including cholesteric liver failure, ischemic reperfusion injury, and neurologic diseases (Gandhi et al., 2009; Izzo et al., 2016; Zheng et al., 2019). MPT-driven cell death was previously referred as regulated necrosis without clear mechanism. Our study establishes that MPT can be sensed by Apaf-1 pyroptsome to trigger caspse-3/GSDME mediated pyroptosis and provides insights into understanding how mitochondrion controls pyroptosis, in addition to previously well-established role in apoptosis, necroptosis, and ferroptosis(Bock and Tait, 2019). Although ANT may not represent indispensable MPTP components(Kokoszka et al., 2004), our results support that ANT1 is an important facilitator in bile acids induced MPT and thereafter pyroptotic cell death. ANT1, as an adenine nucleotide translocator, may facilitate the immediate release of ATP from the mitochondria to cytoplasm that is pivotal for the pyroptosome assembly in the cytoplasm.

Caspase-1 is activated in a molecular platform defined as inflammasome in which diverse PRRs recruit ASC to form a protein complex scaffold. Caspase-9 has been well established to be activated in Apaf-1 adapted apoptosome, while caspase-8 is activated upon death receptors engaging of a protein complex named as death-inducing signaling complex (DISC) using FADD as the adaptor. Here we found that caspase-4/11 can be also recruited and activated in the Apaf-1 protein complex, resulted in a facilitated pyroptosis within 4 hrs instead of apoptosis, in spite of caspase-3 serving as the executioner protease in both cases. Therefore, we term here the MPT triggered assembly of protein complex as Apaf-1 pyroptosome. It seems that the more extensive and earlier activation of caspase-3 triggered by caspase-4 pyroptosome preferably cleaves GSDME to induce pyroptosis, which is an irreversible process and thereby sparing too little time for caspase-3 to cleave the substrates engaged in apoptotic events. Although the exact nature of how cells differentially dictate the choice between apoptosis and pyroptosis warrants extensive future research, our results indicate that pyroptosis necessitates much stronger stimuli and higher threshold (in terms of the rate and amount of cytochrome c and ATP release proposed in this study). Notably, MPT elicited pyroptosis takes place at a much more narrow time window (<4 hrs) than that for MOMP induced apoptosis (usually>24 hrs), and also chemotherapeutic drugs in GSDME positive cells (usually>18 hrs)(Wang et al., 2017). Thus, the assembly of caspase-4 pyroptosome may represent an immediate responsive mechanism for cells detecting extremely dangerous signal for facilitated removal of dying cells.

Caspase-4 can be also auto-activated upon ligation of LPS and thereby cleaving GSDMD for triggering pyroptosis. In contrast, caspase-4 activated in the Apaf-1 pyroptosome preferentially activates caspase-3 and thereby GSDME; GSDMD in this setting is erroneously cleaved to be an inactive form by caspase-3. We provide evidence that GSDMD is needed to be firstly bound with LPS/caspase-4 complex for correct cleavage while GSDMD cannot be recruited to caspase-4 pyroptosome and thus cannot be correctly cleaved. It awaits future research at atom levels to elucidate the exact nature of how caspase-4 is activated in conditions of LPS ligation and in Apaf-1 pyroptosome as that for caspase-9 activation in apoptosome(Hu et al., 2014). Such research will be also important for intensive understanding of how cells dictate the choice between apoptosis and pyroptosis.

Together with previous findings, our results indicate that caspase-4 can detect danger signals (PAMPs and DAMPs) via distinct modes either by direct binding or MPT-triggered Apaf-1 pyroptosome assembly. These findings may shed lights on understanding how pathogen infections meet with host-derived DAMPs in aggravating the pathological processes, and indicate that the mitochondria have evolved a precise machinery in quantitative sensing of stress for culminating appropriate types of cell death. Moreover, the definition of MPT-driven cell death as caspase-4 sensed pyroptosis may provide insights to the development of therapeutics for MPT-driven diseases such as cholesteric liver failure and ischemia tissue damage.

## ACKNOWLEDGEMENTS

This work was financially supported by National Natural Science Foundation of China (grants 81720108032, 81930109 and 81421005 to H. Hao; 81973559 and 81603193 to L.C; 81530098 and 81421005 to G.W.); the Project for Major New Drug Innovation and Development (grants 2018ZX09711001-002-003 to H.Hao; 2018ZX09711002-001-004 to L.C.); Overseas Expertise Introduction Project for Discipline Innovation (G20582017001) to H. Hao; “Double-First Class” initiative project (CPU2018GF09) and Sanming Project of Medicine in Shenzhen (grants SZSM201801060 to H. Hao).

We greatly appreciate Dr. Feng Shao at National Institute of Biological Sciences (Beijing, China) for kindly providing caspase-1^-/-^, *caspase-11*^-/-^, *caspase-1/11*^-/-^, *GSDMD*^-/-^ and *GSDME*^-/-^ mice.

## AUTHOR CONTRIBUTIONS

H. Hao conceived the project; H. Hao and L.C. designed the study; W,X.,Y.C. and L.C. carried out most experiments and collected and analyzed data; Q.Z., H.Huang, C.D. and Y.W. participating some experiments; H. Hao., L.C. and W.X. wrote and revised the manuscript; G.W. was responsible for supervising and major revision; W,X. and Y.C. contributed equally to this work.

## STAR★METHODS

### CONTACT FOR REAGENT AND RESOURCE SHARING

Further information and requests for reagents may be directed to, and will be fulfilled by the Lead Contact Haiping Hao. All unique reagents generated in this study are available from the Lead Contact without restriction or require a completed Materials Transfer Agreement if there is potential for commercial application.

### EXPERIMENTAL MODEL AND SUBJECT DETAILS

#### Animals

Wild-type (WT) C57BL/6J male mice were purchased from Beijing Vital River Laboratory Animal Technology Co., LTD (Beijing, China) at the age of 5 weeks. *Caspase-1^-/-^*, *Caspase-1/11^-/-^, Caspase-11*^-/-^, *GSDMD*^-/-^ and *GSDME*^-/-^ mice on C57BL/6J genetic background were grifts from Prof. Feng Shao at National Institute of Biological Sciences (Beijing, China). All animal experiments were conducted with sex- and age-matched mice and performed with approval from China Pharmaceutical University Animal Care and Use Committee. All mice were kept under controlled conditions of humidity (50±10%), light (12/12 hours light/dark cycle), temperature (25 ± 2°C) and given free access to a standard chow-diet and water ad libitum. All mice were quarantined for one week before starting the experiments.

#### Cell Lines

Human acute monocytic leukemia cell line THP-1 (ATCC, TIB-202) and murine fibroblast cell line L-929 (ATCC, CCL-1) were all kindly obtained from the Stem Cell Bank of the Chinese Academy of Sciences (Shanghai, China). Liver hepatocellular cell line HepG2 (ATCC, HB-8065) and human embryonic kidney cells 293 HEK293T (ATCC, CRL-3216) were purchased from ATCC (Virginia, USA). Cells were maintained in 5% CO_2_ at 37 °C and grown in RPMI-1640 medium for THP-1, DMEM medium for HepG2 and HEK293T, and DME/F-12 medium for L-929 containing 10% Fetal Bovine Serum (FBS), 2 mM L-glutamine, penicillin (50 U/mL) and streptomycin (100 mg/mL) (All medium and supplements were obtained from GIBCO). THP-1 cells were differentiated into macrophages with 10 ng/mL phorbol 12-myristate 13-acetate (PMA, P8139, Sigma-Aldrich) for 48 hrs. All cell culture medium or compound dilutions were decontaminated from endotoxin and mycoplasma by following the instructions of Toxin Sensor− limuloid reagent detection kit (L00350, Genscript) and Mycoplasma Detection Kit (CA1080, Solarbio).

#### Isolation of Bone Marrow-Derived Macrophages

Murine bone marrow-derived macrophages (BMDMs) were harvested from hind legs as previously described(Zhang et al., 2008). The isolated BMDMs were finally resuspended and cultured in the DMEM/F-12 complete medium containing 20% of L-929 conditioned medium, 10% FBS, penicillin (50 U/mL) and streptomycin (100 mg/mL) (All medium and supplements were obtained from GIBCO). The medium was renewed every 3 days and nonadherent cells are eliminated, the cells were used for assays on 7^th^ day.

#### Plasmids and Recombinant Proteins

Human APAF1(1-105AA)-pcDNA3.1(+)-AMP-His (contract No. #190731HE5210-2R3), APAF1(1-591AA)-pcDNA3.1(+)-AMP-His (#190806HE5934-4R8), APAF1-pcDNA3.1(+)-AMP-His (#190623S2HE318-8R26) and CASPASE-9-pcDNA3.1(+)-AMP-Myc (#190403HE9532-5R9) overexpression plasmids were constructed by and obtained from Sangon Biotech (Shanghai, China). Human CASPASE-4-pcDNA3.1(-)-AMP-HA (#MY3892-1), CASPASE-4-pcDNA3.1(-)-AMP-His (#DT3646-1), CASPASE-4(CARD)-pcDNA3.1(-)-AMP-His (#MY3892S) overexpression plasmids were from Detai Bio (Nanjing, China). Recombinant human pro-caspase-4-GFP-His (#DT2148C), recombinant murine pro-caspase-11-GFP-His (#DT2090C) and recombinant human caspase-4 (CARD)-Fc-HA (#MY3897S) were synthetized by and obtained from Detai Bio (Nanjing, China). Recombinant human Apaf-1(1-591AA)-His (#PPL20190730PL02) and recombinant human caspase-9 (CARD)-GST-Myc (#PPL20190730PL02) were synthetized by and obtained from BioGot (Nanjing, China). Other plasmids and recombinant proteins were commercially obtained as stated in Key Resources Table.

### METHOD DETAILS

#### Bile Duct Ligation

All animals were intraperitoneal injected with chloral hydrate solution (0.3 g/kg). After anesthesia, a median abdominal incision was made and the common bile duct was found and carefully ligated with 7–0 Prolene (Ethicon, Somerville, NJ). The bile duct was dissected without ligation in sham operation, then wiped the incision with alcohol swab and injected with 0.5 ml of 0.9% saline on incision to improve recovery and survival after operation. Kept the mice warm at 37.5 °C until recovery. Mice were then used for experiment 7 or 14 days after surgery; survival was monitored every day after surgery until 14^th^ day.

#### Cell Treatment

Unless otherwise indicated in figure legends, cells were treated with 200 μM deoxycholic acid (DCA) or chenodeoxycholic acid (CDCA) for 4 hrs, 20 μM TG for 1hr or 0.5∼1 mM LND for 12 hrs to induce pyroptosis. For LPS induced pyroptosis, cells were primed with 1μg/mL LPS for 12 hrs, then 5 μg/mL of LPS or 3 μg/mL MDP was transfected with 0.3% (v/v) Fugene/HD and 33 uL Opti-MEM in 200 μL medium for 16hrs. In mRNA expression experiments, cells were treated with 200 μM bile acid or 1μg/mL LPS for 4 hrs. To induce apoptosis, cells were treated with 50 μM bile acid for 16 hrs, or 50 ng/mL TNFα/10 μg/mL cycloheximide (CHX), doxorubicin (Dox, 10 μM), 5-flurouracil (5-FU, 20 μM) for 12 hrs.

#### CRISPR/Cas9 Knockout

*APAF-1* (sc-401291), *CASPASE-4* (sc-400858) and *CASPASE-9* (sc-400257-KO-2) CRISPR/Cas9 plasmids were all purchased from Santa Cruz (Dallas, USA). HepG2 was seeded in a 6-well plate at a density of 5 x 10^5^/well and cultured in antibiotic-free DMEM medium for 24 hrs before transfection. 2.5 µg of plasmid and 8 µL of UltraCruz® Transfection Reagent (sc-395739) were premixed by 300 µL plasmid transfection medium (sc-108062) and then brought up to a final volume of 3 mL by DMEM medium. Replace cell culture with 3 mL transfection buffer per well and cells were used for assay at 48 hours after transfection.

#### SiRNA Transfection

*ANT1* (Assay ID: 119895), *ANT2* (Assay ID: 119592) siRNA and negative control siRNA (12935100) were from Invitrogen (Carlsbad, USA). siRNA was transfected into HepG2 cells with Lipofectamine RNAiMAX (13778030, Invitrogene) by using a reverse transfection method according to instructions. Briefly, 500 μL of Opti-MEM medium, 5 μL Lipofecramine RNAiMAX and 20 nM siRNA were premixed per well in a 6-well plate. HepG2 cells were plated at a density of 5×10^5^ per well, optimal transfection time was assessed by the maximal silence efficiency evidenced from western blot analysis, and cells were used for assay 48 hours after transfection.

#### Plasmid Transfection and Pull-down Assay

Plasmid transfection: HEK293T cells were seeded in a 6-well plate at a density of 3 x 10^5^/well and plasmid were transfected into cells with Lipofectamine 3000/P3000 according to instructions. Briefly, 2.5 μg plasmid, 5 μL P3000 and 5 μL Lipofectamine 3000 were mixed into 500 μL Opti-MEM medium in order, then added up to 3 mL with DMEM medium before transferring to 6-well plate. 6 hrs after transfection, replaced the cell culture with DMEM containing 10% FBS, and cells were used for assay 48 hours after transfection.

Pull-down assay: Cells were treated with 200 μM DCA or CDCA for 30 min, collected and lysed cells with NP-40 lysis buffer (P0013F, Beyotime). Lysate with 400 μg protein was diluted into 400 μL by protein binding buffer (50 mM NaH2PO4, 300 mM NaCl, 20 mM imidazole, pH 8.0) and then added with 80 μl Ni-NTA Magnetic Agarose Beads (36111,Qiagen) for His-tag pull-down or Anti-HA Magnetic Agarose Beads (B26201, Bimake) for HA-tag pull-down. Slightly rotated the tubes for 1 hr at 4 °C, separate and discard the supernatant with a magnetic separator, wash the beads twice with wash buffer (for Ni-NTA beads : 50 mM NaH2PO4, 300 mM NaCl, 20 mM imidazole, 0.005% (v/v) Tween-20, pH 8.0; for Anti-HA beads : PBS containing 0.005% (v/v) Tween-20, pH 7.4) and resuspended in 100 μL 2×loading buffer, immediately boiled at 96 °C for 15 min for western blot analysis.

#### Cell Viability and Cytotoxicity Assays

1 x 10^5^ cells /well in a 96-well plate were treated as indicated; cells were used to cell viability determination by using Cell Counting Kit-8 (DJDB4000X, Dojindo Molecular Technologies) and supernatant were collected for cytotoxicity analysis by Lactic Dehydrogenase Release Assay Kit (C0016, Beyotime) according to instructions, respectively.

#### Cytokine Measurement

Human and mouse IL-1α and IL-1β ELISA Kits were all purchased from Excell Bio (Shanghai, China), mouse HMGB1 ELISA kit was from CUSABIO (Wuhan, China). Cell were treated as indicated, collected the supernatant and diluted to appropriate concentration for assay according to instructions. In animal experiments, serum was used for detection of cytokine levels.

#### Recombinant Protein Incubation Assays

Direct caspase-4 activation assessment: Bile acids or LPS were incubated with 250 μM recombinant murine caspase-11-eGFP-His or 100 μM recombinant human caspase-4-eGFP-His protein in a 100 μl reaction system (reaction buffer: 50 mM HEPES (pH 7.5), 150 mM NaCl, 3 mM EDTA and 0.005% (v/v) Tween-20 and 10 mM DTT) at 37 °C for 30 min or 4 hrs. Related to Figure S3B and S3C.

Apaf-1 pyroptosome/apoptosome complex assembling: 400 nM Apaf-1was preincubated with 200 nM cytochrome C and 5 mM ATP overnight at 4 °C in a 100 μL reaction system (reaction buffer: 100 mM KCl, 20 mM HEPES (pH 7.5) and 5 mM DTT). Then, 1 μM pro-caspase-4-eGFP-His was added and incubated for 2 hrs at 37 °C, followed by addition of 200 nM caspase-3 with or without 200 nM GSDMD, and incubated at 37 °C for 1 hr. Parallelly, 1 mg/mL LPS was directly incubated with 1 μM pro-caspase-4-eGFP-His for 2 hrs at 37 °C, 200 nM caspase-3 and GSDMD were then added to reaction system for another 1 hr. 2 U active-caspase-4 incubated with caspase-3 for 1 hr at 37 °C were included as a positive control. Related to Figure 5H-J.

For enzymatic characterization of caspase-4/-9: 700 nM Apaf-1(1-591) was pre-incubated with 5 mM ATP overnight at 4 °C, then incubated with increasing concentrations of caspases (0.025, 0.05, 0.1, 0.2, 0.3, 0.4, 0.5, 0.6 μM) for 2 hrs at 37 °C followed by caspase activity detection. Related to Figure 5L.

For Size-Exclusion chromatography analysis (SEC): 4 μM Apaf-1(1-591)-His was pre-incubated with 5 mM ATP overnight, then added with 2 μM caspase-4-CARD-Fc-HA or caspase-9-CARD-GST-Myc and incubated for 2 hrs at 37 °C in a reaction system of 100 μL. Related to Figure 5M.

For calculation of Kcat/Km of caspase-4/-9: 700 nM Apaf-1(1-591) was pre-incubated with 5 mM ATP overnight at 4 °C, then incubated with 400 nM caspase-9 or 200 nM caspase-4 for 2 hrs at 37 °C. Related to Figure S5J and S5K.

For assessment of ATP threshold in activation of pyroptosome/apoptosme: 400 nM Apaf-1(1-591) was incubated with 0, 0.0625, 0.125, 0.25, 0.5, 1, 2, 4 or 8 mM ATP overnight at 4 °C, then 200 nM caspase-4 or −9 was added respectively or together into reaction system and incubated at 37 °C for 2 hrs. Activation of caspase was detected by immunoblotting. Related to Figure 6J.

#### Caspase Activity Assays

Direct caspase-4 activation assessment: Caspase-4/11 activity was detected by a fluorogenic caspase substrate Z-VAD-AMC. Z-VAD-AMC was added into the reaction system at a final concentration of 75 mM. The reaction mixture was transferred to a 96-well plate and incubated at 37°C for 30 min. Substrate cleavage was monitored by measuring the fluoresce detected at ex365 nm/em 450 nm on a Synergy H1 Microplate Reader (Bio Tek, Winooski, USA).

For cell lysates samples: caspase activities in bile acids, doxorubicin or LPS transfection treated cells as indicated were determined following the instructions of Caspase Colorimetric Assay Kit (caspase-1, K111; caspase-3, K106; caspase-4, K127; caspase-9, K119, Biovision), and detected by a Synergy H1 Microplate Reader (Bio Tek, Winooski, USA). Caspase activities were indicated as relative folds change to control.

For caspase activity assays in pyroptosome assembling incubation buffer: caspase substrate (Ac-LEVD-pNA for caspase-4, Ac-LEHD-pNA for caspase-9 and Ac-DEVD-pNA for caspase-3) was added into reaction buffer to a final concentration of 200 µM, and incubated at 37 °C for 2 hrs. Read samples at 405 nm on a Synergy H1 Microplate Reader (Bio Tek, Winooski, USA). Caspase activities were indicated as relative fold changes to caspase control.

#### Microscale thermophoresis (MST) assay

Analyses of direct binding of between molecules were performed by use of Monolith™ NT.115 instrument (Nano Temper Technology. Munich, Germany). Recombinant pro-caspase-4/11-eGFP-His were desalted and fluorescent labeled in advance by use of a Monolith™ NT Protein Labeling Kit RED-MALEIMIDE (MO-L004, Nano Temper Technology, Munich, Germany) according to instructions. For measurement binding between bile acid and caspase-4/11: Gradient concentrations of bile acid (0.01, 0.1, 1, 10, 100 and 1000 μM) and LPS (31.25, 62.5, 125, 250, 500, 1000 and 2000 nM) were incubated with 0.2 μM pre-labelled recombinant pro-caspase-4/11-eGFP-His at a volume ratio of 1:1 for 30 min at room temperature. For measurements of the binding between Apaf-1 and caspase-4: recombinant human Apaf-1 was preincubated with Cytochrome C (Cyt-C, half concentrations of Apaf-1, 709-CCH-010, R & D System) and 5 mM ATP (A6559, Sigma-Aldrich) overnight at 4 °C to activate Apaf-1, then gradient concentrations of recombinant human Apaf-1 (23, 46, 92, 184, 370 and 740 nM) were incubated with 1 μM recombinant pro-caspase-4-eGFP-His for 30 min at room temperature.

10 μL of incubated mixture was inhaled into capillary and loaded onto Monolith™ NT.115 instrument, then perform a cap scan of all the capillaries and run the MST experiment using 40% LED-power and 80% MST-power. Data were analyzed using the NT Analyses 1.5.41 software. eGFP-His protein was conducted parallelly as a negative control.

#### Flow Cytometry

For flow cytometry analysis, 2 x 10^6^ cells of each group in 6-well plate were collected with 0.25% typsin without EDTA (40101ES25, Yeasen Biotech, Shanghai, China), centrifuged at 2500 rpm for 5 min and washed pellet once with PBS, cells were resuspended with 200 µL 1 x binding buffer and stained with propidium iodide (556547, BD Pharmingen, San Diego, USA) for 15-20 min at room temperature. 1 x 10^4^ cells were collected and analyzed by BD Accuri C6 plus flow cytometer (BD Biosciences, San Jose, USA).

#### Mitochondrial Membrane Potential Assay

The change of mitochondrial membrane potential was assayed by using JC-1 mitochondrial membrane potential assay kit (C2006, Beyotime Biotechnology). In brief, 1x 10^5^/well BMDMs were seeded into a 96-well plate and treated as indicated in figure legends. Replaced culture medium with 100 µL JC-1 probe solution and incubation/wash according to instructions. JC-1 were detetect by a fluorescent reader (Synergy H1 Microplate Reader, Bio Tek, Winooski, USA) under Ex490/Em530 nm for monomer and Ex525/Em590 nm for polymer (aggregate). The ratio of aggregates/monomers suggested the dissipation of mitochondrial transmembrane potential and data were relative to vehicle/control. CCCP was applied as a positive control inducing JC-1 depolymerization.

#### Mitochondrial Permeability Transition Pore (MPTP) Detection

Cells were seeded into a 96-well plate at a density of 1x 10^5^/well and treated as indicated in figure legends. MPTP opening was determined following the instructions of Mitochondrial Permeability Transition Pore Assay Kit (K239-100, Biovision) with minor changes. Briefly, diluted calcein-AM solution to 500 nM with solution buffer, washed the cells once with wash buffer and incubated cells with 50 µL calcein-AM solution for 20 min at 37 °C, then added with 50 µL CoCl_2_ solution and incubated for another 10 min to rule out excessive calcien-AM, brought the volume up to 100 µL. Washed with wash buffer thrice and read the plate under Ex490/Em515 nm with fluorescent reader (Synergy H1 Microplate Reader, Bio Tek, Winooski, USA). The decline rate of fluorescent value indicated the degree of MPTP opening and the percent of each group was relative to vehicle/control. 200 µM arachidonic acid (AAs) was used as a positive control.

#### Microscopy Imaging of Cell Death and Immunofluorescence

To observe cell morphology changes, 3 x 10^4^ BMDMs or differentiated THP-1 cells were plated in cover-glass bottom dishes and treated as indicated in figure legends. Cells were washed twice with PBS and incubated with 200 µL 1x binding buffer containing 2.5 µL propidium iodide (556547, BD Pharmingen, San Diego, USA) for 15-20 min at room temperature. Washed the cells thrice with PBS. Cell images were acquired on LSM700 confocal microscope (Zeiss, Oberkochen, Germany).

For colocalization analysis of caspase-4 and APAF-1, cells were washed twice with PBS/T after treatment with 200 μ bile acids for 4 hrs or 1mM LND for 6 hrs, fixed with ice-cold 4% paraformaldehyde for 20 min, permeabilized using 0.2% Triton X-100 in 3% BSA for 15 min, blocked with 3% BSA for 1 hr at room temperature. Then cells were incubated with primary antibodies of caspase-4 (1:50, ab25898, Abcam) and Apaf-1 (1:50, sc-65891, Santa Cruz) at 3% BSA overnight at 4°C.

Washed cells six times with PBS/T (PBS containing Tween-20, 0.05% v/v) before incubating with Donkey Anti-Rabbit IgG(H+L) Antibody, Alexa Fluor 488 (1:1000, A-21206, Invitrogen) and Donkey Anti-Mouse IgG (H+L) Antibody, Alexa Fluor 555 (1:1000, A-31570, Invitrogen) for 1 hr at 37 °C in 3% BSA, then washed cells five times with PBS/T. Cell images were acquired on LSM700 confocal microscope (Zeiss, Oberkochen, Germany).

#### Electron Microscopy

Cell or murine liver samples were prepared as previously described with minor changes(Kravchenko et al., 2006). 1 x 10^6^ BMDMs treated as indicated or 1 mm^3^ of liver section from mice were fixed in 0.2 M phosphate buffer (KH2PO4/Na2HPO4, pH 7.5) supplemented with 2.5% glutaraldehyde (G5882, Sigma-Aldrich) for 2-4 hrs. Washed with phosphate buffer 15 min for thrice and samples were post-fixed for 60 min in 1% osmium tetroxide (75632, Sigma-Aldrich). After washed in phosphate buffer, samples were dehydrated in gradient ethanol solutions (50%, 70%, 80%, 90%, 95% and 100%, 15 min each step). The samples were then infiltrated sequentially in 1:1 (v/v) of ethanol: spurr resin (Polyscience, Warrington, USA) and 1:3 of ethanol: spurr resin for 30 min respectively, 100% spurr resin for 3 hrs and finally 100% spurr resin for 24 hrs at 60°C for polymerization. Ultra-thin sections were isolated on nickel grids and stained for 10 min in 2% uranyl acetate and then in reynolds lead citrate for 15 min. Examined at 80 kV using a Hitachi H7650 transmission electron microscope (Tokyo, Japan).

Cristae density was calculated by using Image pro for area void of cristae as previously described(Luongo et al., 2017) and mitochondrial size was calculated by tracing individual mitochondrion after calibration for distance (minimum of 40 mitochondria were included per mouse/group, n = 3).

#### Size-Exclusion Chromatography (SEC) Analysis

Sample preparation : To further determine the stoichiometric ratio of Apaf-1 and caspase-4 in the pyroptosome scaffold, constructed Apaf-1(1-591AA)-His/caspase-4-CARD-Fc-HA complex was isolated from incubation system via double purification by HA-tag and His-Tag pulldown as described in pull-down assay. Then, the stable complex was identified by under non-denaturing condition, and composition ratio of Apaf-1 and caspase-4 was analyzed by subjecting to SEC after denaturing with 70% acetonitrile. Apaf-1(1-591AA)-His/caspase-9(CARD)-GST-Myc complex was constructed and parallelly analyzed as a control.

SEC condition: Advance Bio SEC 300A 2.7 μm 7.8 x 150 mm (PL1180-3301, Agilent Techology) column was used in this study. The column was preequilibrated with 50 mM Na_2_HPO_4_·12H_2_O, 200 mM NaCl, pH 7.5 and calibrated with molecular weight standards (69385, Sigma). Sample was injected and eluted with a flow rate of 0.5 ml/min.

Calculation: On the basis that the peak areas proportions that correspond to caspase-4 (CARD) and Apaf-1 (1-591AA) were constant at OD 280 nm, the response ratio between caspase-4 (CARD) and Apaf-1 (1-591AA) was calculated by series equivalent concentrations of caspse-4 (CARD) and Apaf-1 (1-591AA). Composition ratio of Apaf-1 and caspase-4 was quantified by relative peak areas normalized by response ratio.

#### ATP quantification assay

1 x 10^7^ BMDMs were treated as indicated. Collected cells with 0.25 % Typsin-EDTA (25200072, Gibco, NY, USA) and centrifuged at 2500 rpm for 5 min. Cytoplasm and mitochondrion were isolated with Cell Mitochondria Isolation Kit (C3601, Beyotime Biotechnology, Shanghai, China), 100 µM ATPase inhibitor ARL67156 (A265, Sigma-Aldrich) was used for preventing ATP degradation. ATP Determination Kit (A22066, Invitrogen, Carlsbad, USA) was used for ATP quantification in cytoplasm and mitochondrion by following the instructions.

#### Quantitative Real-Time PCR (qPCR)

qPCR analysis was conducted as previously described(Hao et al., 2017). RNA was extracted using RNAiso Plus reagent (Takara, Japan) according to the manufacturer’s instructions. cDNA was synthesized from 2.5 μg total RNA using Superscript II reverse transcriptase (RR047A, Takara, Japan). Sequences of qPCR primers were shown in the Table S1, mRNA expression levels were normalized by *GAPDH* mRNA levels and represented as fold change relative to control/vehicle group.

#### Co-immunoprecipitation

1×10^7^ BMDMs cells, or 20 mg liver tissue homogenate were harvested and lysed in 300 μL NP-40 lysis buffer containing protease inhibitor cocktail (P8340, Sigma-Aldrich). Lysates with 200 μg protein were diluted into 300∼400 μL and used for co-immunoprecipitation analysis according to instructions. Briefly, lysates were pre-cleared with Protein A Agarose beads (15918-014, Invitrogen) for 1.5 hrs at 4°C, and antibody-protein A Agarose beads was pre-constructed by incubation of 2μg Apaf-1(8969, Cell Signaling Technology) or 4 μg caspase-11(180673, Abcam) antibody with 50 μL Protein A Agarose overnight at 4 °C in PBS. Then, incubated lysates with antibody-protein A Agarose beads for 3 hrs at 4°C. Wash with NP-40 lysis buffer for three times, the immunoprecipitates were boiled up for 10 min in 80 μl of 2× loading buffer before western blot analysis. All beads were pre-blocked by 5% BSA for 1hr at room temperature to exclude nonspecific binding before use. Cell or tissue lysates containing 60 μg protein were used to western blot analysis as input.

#### Western blotting

The collected cells or liver tissue homogenates were lysed in RIPA lysis solution containing 1x protein inhibitor cocktail (P8340, Sigma-Aldrich) and quantitated by BCA assay (all protein extraction and quantification reagents were from Beyotime Biotechnology, Shanghai, China). Dilute the lysates with XT Sample Buffer (1610791, Bio-Rad) and boiled up for 5 min, 60 µg protein was then subjected to electrophoresis by use of Mini-Protean Tetra System (Bio-Rad, Hercules, USA), transferred to PVDF membrane (1620177, Bio-Rad) by use of a Trans-Blot Turbo Transfer System (Bio-Rad, Hercules, USA) and probed with primary antibodies followed by HRP-conjugated secondary antibodies. The immunoreactive bands were visualized with HRP substrate (170-5061, Bio-Rad) and detected by using an iBright CL1000 System (Invitrogen, Carlsbad, USA). Blots were analyzed by iBright Analysis Software Version 3.0.1 (Invitrogen, Carlsbad, USA).

For the detection of cytochrome c release in cytoplasm, cytoplasmic proteins were isolated following the instructions of Cytochrome c Releasing Apoptosis Assay Kit (K257-100, Biovision). Cytoplasmic protein extractions were diluted with XT Sample Buffer (1610791, Bio-Rad) and boiled up for 5 min before western blot analysis. Antibody of cytochrome c was provided by kit.

The following antibodies were used in immunobloting: GSDME (ab215191, Abcam); ANT1 (ab110322, Abcam); HA (ab18181, Abcam); Myc (ab9132, Abcam); GAPDH (AB0037, Abways Technology); Caspase-3 (9662, Cell Signaling Technology); Caspase-9 (9508, Cell Signaling Technology); PARP (9532, Cell Signaling Technology); ANT2 (14671, Cell Signaling Technology); His (12698, Cell Signaling Technology); Apaf-1 (sc-65891, Santa Cruz Biotechnology); Caspase-1 (sc-56036, Santa Cruz Biotechnology); Caspase-4 (for sc-56056, Santa Cruz Biotechnology); HMGB1 (BS1918, Bioworld Technology); GSDMD (NBP2-33422, Novus); GSDMD (sc-393656, Santa Cruz Biotechnology); HRP Conjugated Anti-Rabbit IgG (H+L) antibody (ab6721, Abcam); HRP Conjugated Anti-Mouse IgG (H+L) antibody (ab6789, Abcam) and HRP Conjugated Anti-Goat IgG (H+L) antibody (ab6885, Abcam).

### QUANTIFICATION AND STATISTICAL ANALYSIS

Data are presented as mean ± SEM, number of replicates is stated in the figure legends. At least two independent experiments were performed throughout in this study. For image data, one representative image of three independent experiments is shown. Animal studies were age and sex matched and randomized by bodyweight. The researchers were not blinded and no statistical methods were used to predetermine sample size. Statistical significance between two groups was determined by two-tailed Student’s t-test. For three or more groups, statistical significance was determined by one-way ANOVA analysis. Log-rank (Mantel-Cox) test was used for survival analysis (n=10). p values of < 0.05 were considered significant, otherwise indicated as nonsignificant (n.s.). Graphs and all statistical analysis were performed in GraphPad Prism 6.0 software. Fow cytometry analysis were carried out using BD Accuri C6 plus. Image pro was used to calculate the area void of cristae (%), the area of mitochondria and cristae were traced respectively, area void of cristae (%) = (area of mitochondria-area of cristae) / area of mitochondria, minimum of 40 mitochondria were analyzed per group (n=3). Calculation of dissociation constant Kd were carried out using NT Analyses 1.5.41 software, performed nonlinear curve-fitting of the binding and Kd was calculated automatically.

### DATA AND SOFTWARE AVAILABILITY

All software used is available online, either freely or from a commercial supplier and is summarized in the Key Resources Table. Data will be accessible upon publication of the manuscript. Raw data of immunoblotting analysis can be accessed via doi: 10.17632/hj6bwp284s.1 (http://dx.doi.org/10.17632/hj6bwp284s.1).

## SUPPLEMENTARY INFORMATION LEGENDS

**Figure S1. Bile Acids Activate Caspase-4 Noncanonical Inflammasome and Induce Pyroptosis in THP-1 Cells. (Related to Figure 1)**

**(A)** Representative immunoblots of caspase-1, −4, cleaved (Cl-)caspase-1, −4 and GAPDH in cell lysates (Lys) andr supernatant (Sup) of THP-1 cells, cells were stimulated with 200 μM bile acids for 4 hrs (Cholic acid, CA; Ursodeoxycholic acid, UDCA; Deoxycholic acid, DCA; Chenodeoxycholic acid, CDCA; β-muricholic acid, β-MCA).

**(B)** *CASPASE-4* mRNA levels in THP-1 cells treated with 50 ng/ml LPS or 200 μM BAs for 4hrs.

**(C and D)** CCK8 assay for cell viability **(C)** and LDH release for cytotoxicity **(D)** of BMDMs challenged with 50-400 μM bile acids for 4 hrs (tauro-chenodeoxycholic acid, TCDCA; tauro-deoxycholic acid, TDCA).

**(E)** Representative confocal microscopy analysis of propidium iodide (PI) positive THP-1 cells, scale bars, 50 μm. Cell treatment as in **Figure 1A-D**.

**(F and G)** CCK8 assay for cell viability **(F)** andLDH release for cytotoxicity **(G)** of BMDMs stimulated with 100-800 μM tauro-deoxycholic acid (TDCA) or tauro-chenodeoxycholic acid (TCDCA) for 4 hrs.

**(H)** Representative images of electron microscopy analysis of cell morphology changes in BMDMs. Cells were stimulated with 200 μM bile acids for 4 hrs, or LPS-primed cells were transfected with 5 μg/ml LPS by 0.3% Fugene HD for 16 hrs. Scale bars, 1 μm.

**(I)** Representative confocal microscopy analysis of propidium iodide (PI) positive BMDMs, scale bars, 50 μm. Cell treatment as in (H).

**(J-M)** CCK8 assay for cell viability **(J)**, LDH release for cytotoxicity **(K)** and relative caspase activities **(L and M)** in BMDMs. Cells were treated with 0.002% (v/v) SDS or 200 μM DCA for 4 hrs.

**(N and O)** CCK8 assay for cell viability **(N)** and LDH release for cytotoxicity **(O)** of THP-1 cells; cells were pre-treated with 20 μM RIP1 inhibitor Necrostatin-1 or MLKL inhibitor Necrosulfonamide, following treatment with 200 μM bile acid or 10 ng/mL TNF-α + 10 μM Z-VAD-FMK for 4 hrs. TNF-α/ Z-VAD-FMK was used as a positive control to induce necrosis.

GAPDH was used as internal standard/loading control in qPCR/immunoblot analyses. Bar graphs expressed as mean ± SEM (n=3). ***, *p*<0.001; **, *p* < 0.01; *, *p* < 0.05 compared to vehicle unless indicated otherwise in graphs.

**Figure S2. Bile Acids Induced Pyroptosis Depends on GSDME in HepG2 Cells. (Related to Figure 2)**

**(A and B)** CCK8 assay for cell viability **(A)** and LDH release for cytotoxicity **(B)** of *GSDME* silenced HepG2 cells compared to negative control (N.C.) group; Cells were stimulated with μM DCA or CDCA for 4 hrs.

**(C)** Flow cytometry analysis of Annexin V-FITC/PI staining in N.C. and *GSDME* silenced HepG2 cells, cells were treated as in (A and B).

**(D)** Proportions of PI positive cells, quantified from images as shown in (C).

Bar graphs expressed as mean ± SEM (n=3). *, *p*<0.05; **, *p*<0.01.

**Figure S3. Bile Acids Cannot Directly Bind and Activate Caspase-4/11. (Related to Figure 3)**

**(A)** Microscale thermophoresis (MST) measurements of the binding of caspase-4/11 with bile acids or LPS. Gradient concentrations of bile acid (0.01, 0.1, 1, 10, 100 and 1000 μM, from bottom to top) and LPS (31.25, 62.5, 125, 250, 500, 1000 and 2000 nM, from bottom to top) were incubated with 0.2 μM pre-labelled recombinant pro-caspase-4/11-eGFP-His at a volume ratio of 1:1 for 30 min at room temperature. The calculated dissociation constants (*Kd*) were listed.

**(B and C)** Recombinant casaspe-4, −11 activities were determined by a fluorescent casaspe substrate Z-VAD-AMC. 500 μM bile acid or 1 mg/ml LPS were incubated with 250 nM caspase-11 or 100 nM caspase-4 for 30 min at room temperature.

**(D and E)** CCK8 assay for cell viability **(D)** and LDH release for cytotoxicity **(E)** of BMDMs. Cells were pre-treated with 5 μM Cyclosporin A (CsA), 20 μM N-Ethylmaleimide (NEM) or BKA for 2 hrs followed by 200 μM DCA/CDCA treatment for 4 hrs.

**(F-I)** Calcien-AM/CoCl_2_ assay for MPT **(F)**, LDH release for cytotoxicity **(G)** and IL-1α/β secretion **(H and I)** of BMDMs, LPS-primed BMDMs were pre-treated with were pre-treated with 5 μM CsA, 20 μM NEM or BKA for 2 hrs followed by LPS/Fugene HD transfection for 16 hrs, MPT analysis were carried out at 4 hrs after LPS/Fugene HD transfection.

Bar graphs expressed as mean ±SEM (n=3). **, *p*<0.01; ***, *p*<0.001; n.s., non-significant versus vehicle or between indicated groups in graphs.

**Figure S4. Caspase-4/11 Elicited Pyroptosis Dominants in MPT-Triggered Cell Death. (Related to Figure 4)**

BMDMs were treated with 20 μM TG or 1 mM LND for indicated times in (A-C, F-H), for 1 hr in (D and E) or for 6 hrs in (I and J), 20 μM BKA pretreatment was 2 hrs prior to TG/LND in (D, E, I, J).

**(A, D, F, I)** Δψ_m_ dissipation indicated by JC-1 disaggregation

**(B, E, G, J)** MPT was detected by calcien-AM/CoCl_2_ assay.

**(C and H)** Fold change of cytoplasmic ATP concentration.

Line and bar graphs expressed as mean ±SEM (n=3). **, *p*<0.01; ***, *p*<0.001.

**Figure S5. Both MPT and Apaf-1 Is Necessary for Pyroptosome Assembly.(Related to Figure 5)**

**(A-D)** Calcien-AM/CoCl_2_ assay for MPT **(A)**, CCK-8 assay for cell viability (B), LDH release for cytotoxicity **(C)** and representative immunoblots of caspase-4 and GAPDH in lysates **(D)**. AM/CoCl_2_ assay. HepG2 cells were stimulated with TNFα/CHX (50 ng/mL TNFα + 10 μg/mL CHX), 10 μM Dox or 20 μM 5-FU for 3 hrs in (A) or 12 hrs in (B-D).

**(E)** Representative images by confocal microscopy analysis of the colocalization between Apaf-1 and caspase-4 in HepG2 cells that were stimulated with 200 μM bile acids for 4 hrs or 1 mM LND for 12 hrs. Scale bars, 10 μm.

**(F-H)** CCK-8 assay for cell viability **(F)**, LDH release for cytotoxicity **(G)** and representative immunoblots of Apaf-1, caspase-3, −4, GSDME and GAPDH in lysates **(H)** of *APAF-1* knockout (KO) HepG2 cells compared to WT group; cells were treated with 1 mM LND for 12 hrs.

**(I)** Co-immunoprecipitation analysis of Apaf-1 with caspase-11 in BMDMs lysates. BMDMs were pre-treated with 20 μM BKA for 2 hrs followed by 200 μM bile acid for 1 hr or 1 mM LND for 4 hrs.

**(J and K)** The Michaelis-Menten curve and Kcat/Km of caspase-4 or caspase-9, enzymatic activities of caspase-4 and −9 were detected by substrate Ac-LEVD-pNA and Ac-LEHD-pNA respectively. Incubation systems were constructed as Apaf-1: caspase-4=400 nM: 100 nM and Apaf-1: caspase-9=400 nM: 200 nM.

GAPDH was used as loading control in immunoblot analyses of cell lysates. Bar graphs expressed as mean ±SEM (n=3). *, *p*<0.05; **, *p*<0.01; n.s., non-significant versus control or between indicated groups. The Michaelis-Menten curve was fitted and the value of V_max_ and K_m_ was calculated by using Graphpad prism 6.0, K_cat_ =V_max_/[E], E=enzyme concentration.

**Figure S6. Caspase-9 Is Not Involved in MPT-Elicited Pyroptosis. (Related to Figure 6)**

**(A and B)** Time course analysis of fold change of caspase-1, −3, −4 and −9 activities in HepG2 cells stimulated with 200 μM bile acid.

**(C)** Immunoblotting validation of CRISPR/Cas9 KO of *CASPASE-4* and *CASPASE-9* efficiency in HepG2 cells.

**(D-G)** Relative caspase-4 and −9 activities (D and E), CCK-8 assay for cell viability **(F)** and LDH release for cytotoxicity **(G)** of *CASPASE-4*/*CASPASE-9* KO HepG2 cells compared to WT group. Cells were treated with 200 μM bile acid for 45 min or LPS-primed HepG2 cells were transfected with 5 μg/ml LPS by 0.3% (v/v) Fugene HD for 10 hrs.

**(H)** Flow cytometry analysis of Annexin V-FITC/PI staining in *CASPASE-4*/*CASPASE-9* KO HepG2 cells compared to WT group, cells were treated as in (D-G).

**(I)** Proportions of PI positive cells, quantified from images as shown in (H).

GAPDH was used as loading control in immunoblot analyses of cell lysates. Line and bar graphs expressed as mean ±SEM (n=3). **, *p*<0.01; ***, *p*<0.001.

**Figure S7. Caspase-11 and GSDME Dependent Pyroptosis Dominants BDL Induced Cholesteric Liver Failure. (Related to Figure 7)**

**(A)** Representative immunoblots of ANT1/2 and GAPDH in liver and kidney of mice 7 days and 14 days after BDL/sham (n=6).

**(B-D)** Serum levels of IL-1α, IL-1β and HMGB1 in mice 7 days and 14 days after BDL/sham (n=6).

GAPDH was used as loading control in immunoblot analyses of tissue lysates, kidney was analyzed as an indicator of physiological level of ANT, lysates from six mice was merged by two-into-one to n=3. Scatter plot with box graphs expressed as mean ± SEM. ***, *p*<0.001.

**Table S1. Sequences of qPCR Primers. (Related to STAR★METHODS)**

