## Supplemental figures and table for "Apaf-1 Pyroptosome Senses Mitochondrial Permeability Transition"

**Inventory of supplementary information:**

**Figure S1. Bile Acids Activate Caspase-4 Noncanonical Inflammasome and Induce Pyroptosis in THP-1 Cells. (Related to Figure 1)**

**Figure S2. Bile Acids Induced Pyroptosis Depends on GSDME in HepG2 Cells. (Related to Figure 2)**

**Figure S3. Bile Acids Cannot Directly Bind and Activate Caspase-4/11. (Related to Figure 3)**

**Figure S4. Caspase-4/11 Elicited Pyroptosis Dominants in MPT-Triggered Cell Death. (Related to Figure 4)**

**Figure S5. Both MPT and Apaf-1 Is Necessary for Pyroptosome Assembly. (Related to Figure 5)**

**Figure S6. Caspase-9 Is Not Involved in MPT-Elicited Pyroptosis. (Related to Figure 6)**

**Figure S7. Caspase-11 and GSDME Dependent Pyroptosis Dominants BDL Induced Cholesteric Liver Failure. (Related to Figure 7)**

**Table S1. Sequences of qPCR Primers. (Related to STAR**★**METHODS)**

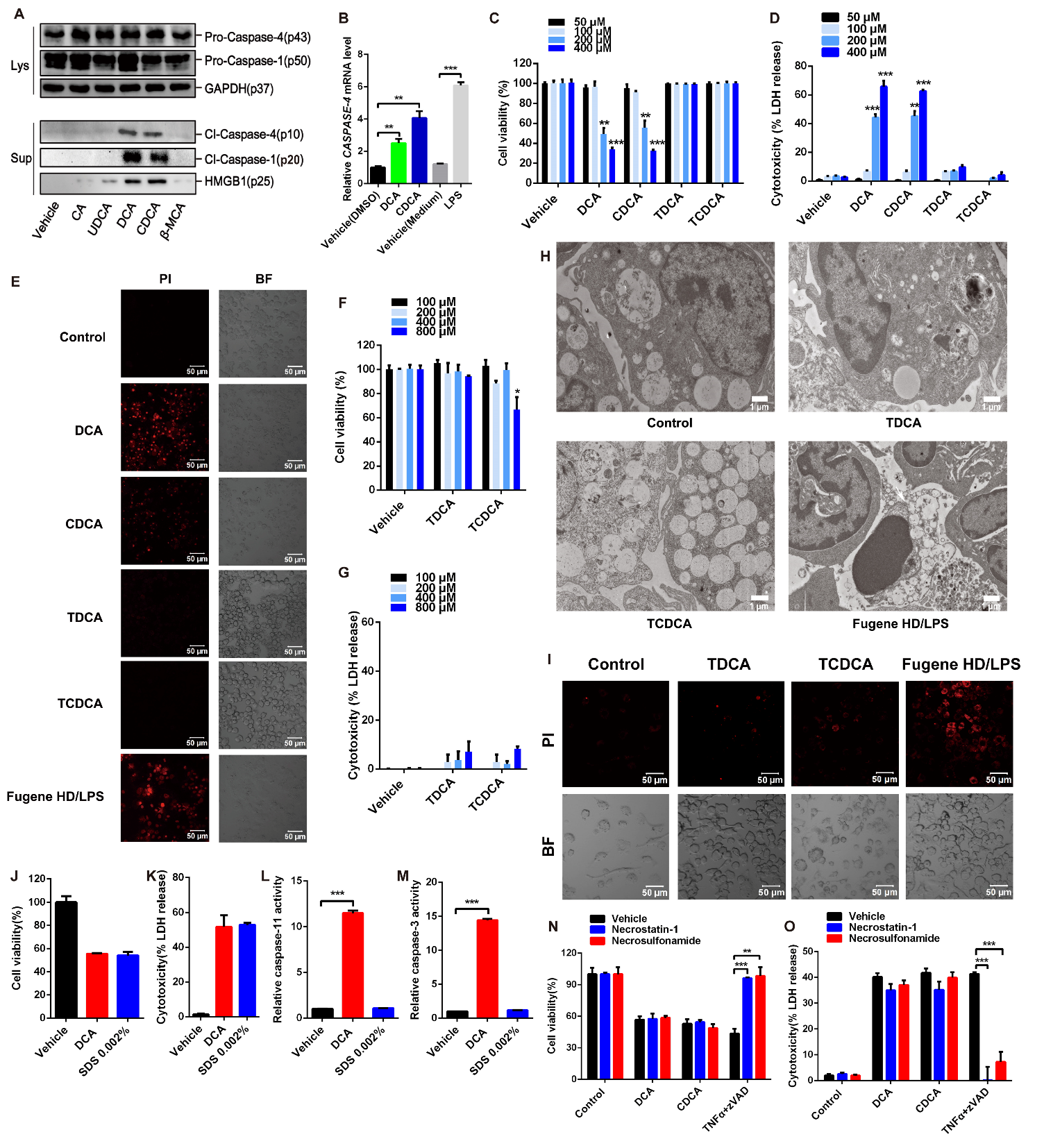

**Figure S1. Bile Acids Activate Caspase-4 Noncanonical Inflammasome and Induce Pyroptosis in THP-1 Cells. (Related to Figure 1)**

**(A)** Representative immunoblots of caspase-1, -4, cleaved (Cl-)caspase-1, -4 and GAPDH in cell lysates (Lys) andr supernatant (Sup) of THP-1 cells, cells were stimulated with 200 μM bile acids for 4 hrs (Cholic acid, CA; Ursodeoxycholic acid, UDCA; Deoxycholic acid, DCA; Chenodeoxycholic acid, CDCA; β-muricholic acid, β-MCA).

**(B)** *CASPASE-4* mRNA levels in THP-1 cells treated with 50 ng/ml LPS or 200 μM BAs for 4hrs.

**(C and D)** CCK8 assay for cell viability **(C)** and LDH release for cytotoxicity **(D)** of BMDMs challenged with 50-400 μM bile acids for 4 hrs (tauro-chenodeoxycholic acid, TCDCA; tauro-deoxycholic acid, TDCA).

**(E)** Representative confocal microscopy analysis of propidium iodide (PI) positive THP-1 cells，scale bars, 50 μm. Cell treatment as in Figure 1A-D.

**(F and G)** CCK8 assay for cell viability **(F)** andLDH release for cytotoxicity **(G)** of BMDMs stimulated with 100-800 μM tauro-deoxycholic acid (TDCA) or tauro-chenodeoxycholic acid (TCDCA) for 4 hrs.

**(H)** Representative images of electron microscopy analysis of cell morphology changes in BMDMs. Cells were stimulated with 200 μM bile acids for 4 hrs, or LPS-primed cells were transfected with 5 μg/ml LPS by 0.3% Fugene HD for 16 hrs. Scale bars, 1 μm.

**(I)** Representative confocal microscopy analysis of PI positive staining BMDMs，scale bars, 50 μm. Cell treatment as in (H).

**(J-M)** CCK8 assay for cell viability **(J)**，LDH release for cytotoxicity **(K)** and relative caspase activities **(L and M)** in BMDMs. Cells were treated with 0.002% (v/v) SDS or 200 μM DCA for 4 hrs.

**(N and O)** CCK8 assay for cell viability **(N)** and LDH release for cytotoxicity **(O)** of THP-1 cells; cells were pre-treated with 20 μM RIP1 inhibitor Necrostatin-1 or MLKL inhibitor Necrosulfonamide, following treatment with 200 μM bile acid or 10 ng/mL TNF-α + 10 μM Z-VAD-FMK for 4 hrs. TNF-α/ Z-VAD-FMK was used as a positive control to induce necrosis.

GAPDH was used as internal standard/loading control in qPCR/immunoblot analyses. Bar graphs expressed as mean ± SEM (n=3). ***, *p*<0.001; **, *p* < 0.01; *, *p* < 0.05 compared to vehicle unless indicated otherwise in graphs.

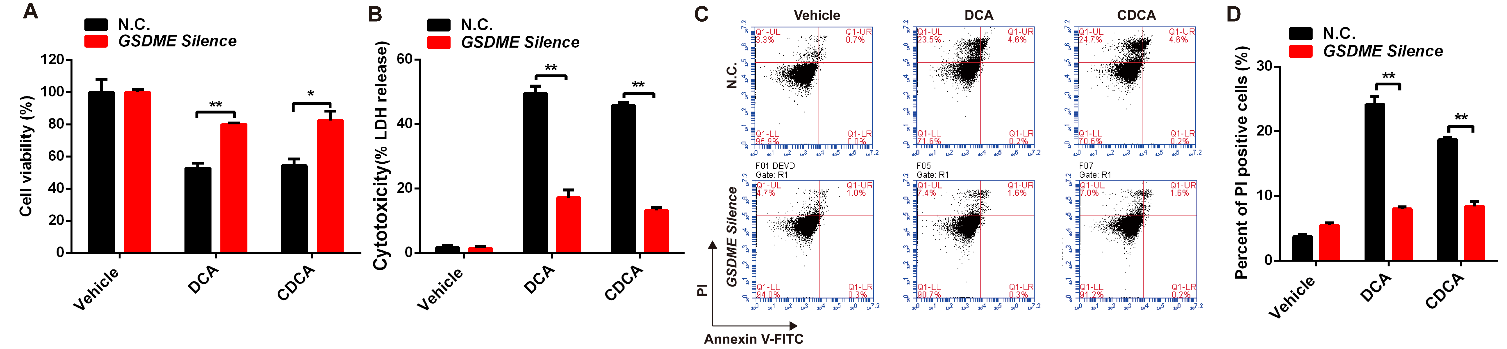

**Figure S2. Bile Acids Induced Pyroptosis Depends on GSDME in HepG2 Cells. (Related to Figure 2)**

**(A and B)** CCK8 assay for cell viability **(A)** and LDH release for cytotoxicity **(B)** of *GSDME* silenced HepG2 cells compared to negative control (N.C.) group; Cells were stimulated with 200 μM DCA or CDCA for 4 hrs.

**(C)** Flow cytometry analysis of Annexin V-FITC/PI staining in N.C. and *GSDME* silenced HepG2 cells, cells were treated as in (A and B).

**(D)** Proportions of PI positive cells, quantified from images as shown in (C).

Bar graphs expressed as mean ± SEM (n=3). *, *p*<0.05; **, *p*<0.01.

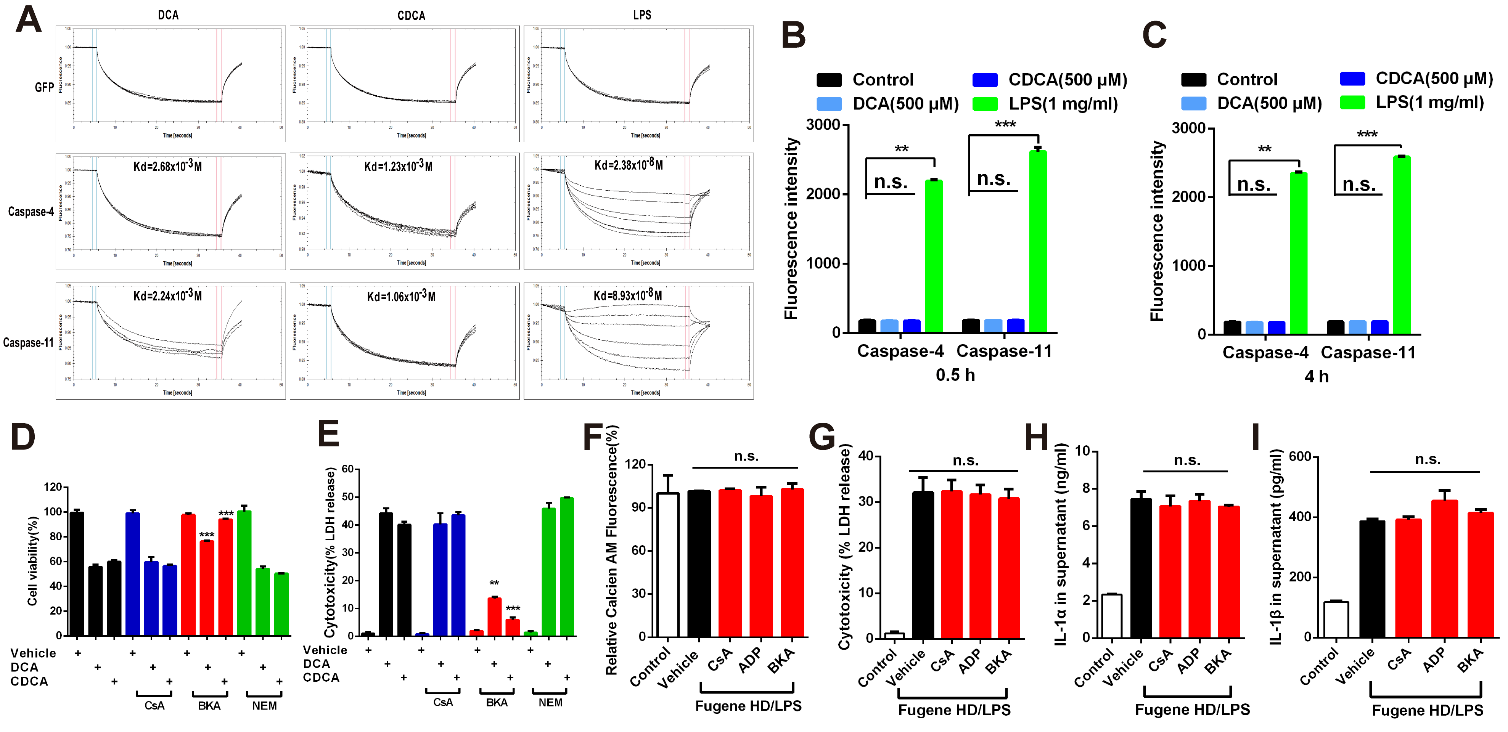

**Figure S3. Bile Acids Cannot Directly Bind and Activate Caspase-4/11. (Related to Figure 3)**

**(A)** Microscale thermophoresis (MST) measurements of the binding of caspase-4/11 with bile acids or LPS. Gradient concentrations of bile acid (0.01, 0.1, 1, 10, 100 and 1000 μM, from bottom to top) and LPS (31.25, 62.5, 125, 250, 500, 1000 and 2000 nM, from bottom to top) were incubated with 0.2 μM pre-labelled recombinant pro-caspase-4/11-eGFP-His at a volume ratio of 1:1 for 30 min at room temperature. The calculated dissociation constants (*Kd*) were listed.

**(B and C)** Recombinant casaspe-4, -11 activities were determined by a fluorescent casaspe substrate Z-VAD-AMC. 500 μM bile acid or 1 mg/ml LPS were incubated with 250 nM caspase-11 or 100 nM caspase-4 for 30 min at room temperature.

**(D and E)** CCK8 assay for cell viability **(D)** and LDH release for cytotoxicity **(E)** of BMDMs. Cells were pre-treated with 5 μM Cyclosporin A (CsA), 20 μM N-Ethylmaleimide (NEM) or BKA for 2 hrs followed by 200 μM DCA/CDCA treatment for 4 hrs.

**(F-I)** Calcien-AM/CoCl_2_ assay for MPT **(F)**，LDH release for cytotoxicity **(G)** and IL-1α/β secretion **(H and I)** of BMDMs, LPS-primed BMDMs were pre-treated with were pre-treated with 5 μM CsA, 20 μM NEM or BKA for 2 hrs followed by LPS/Fugene HD transfection for 16 hrs, MPT analysis were carried out at 4 hrs after LPS/Fugene HD transfection.

Bar graphs expressed as mean ±SEM (n=3). **, *p*<0.01; ***, *p*<0.001; n.s., non-significant versus vehicle or between indicated groups in graphs.

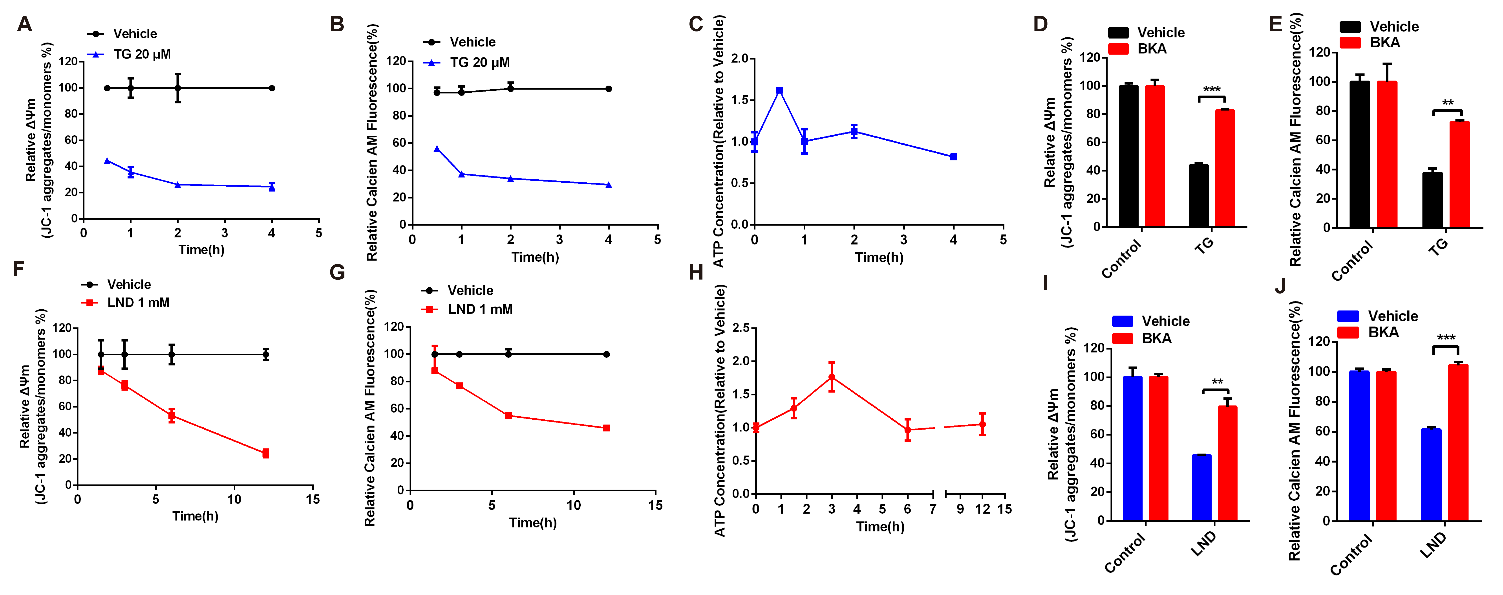

**Figure S4. Caspase-4/11 Elicited Pyroptosis Dominants in MPT-Triggered Cell Death. (Related to Figure 4)**

BMDMs were treated with 20 μM TG or 1 mM LND for indicated times in (A-C, F-H), for 1 hr in (D and E) or for 6 hrs in (I and J), 20 μM BKA pretreatment was 2 hrs prior to TG/LND in (D, E, I, J).

**(A, D, F, I)** Δψ_m_ dissipation indicated by JC-1 disaggregation

**(B, E, G, J)** MPT was detected by calcien-AM/CoCl_2_ assay.

**(C and H)** Fold change of cytoplasmic ATP concentration.

Line and bar graphs expressed as mean ±SEM (n=3). **, *p*<0.01; ***, *p*<0.001.

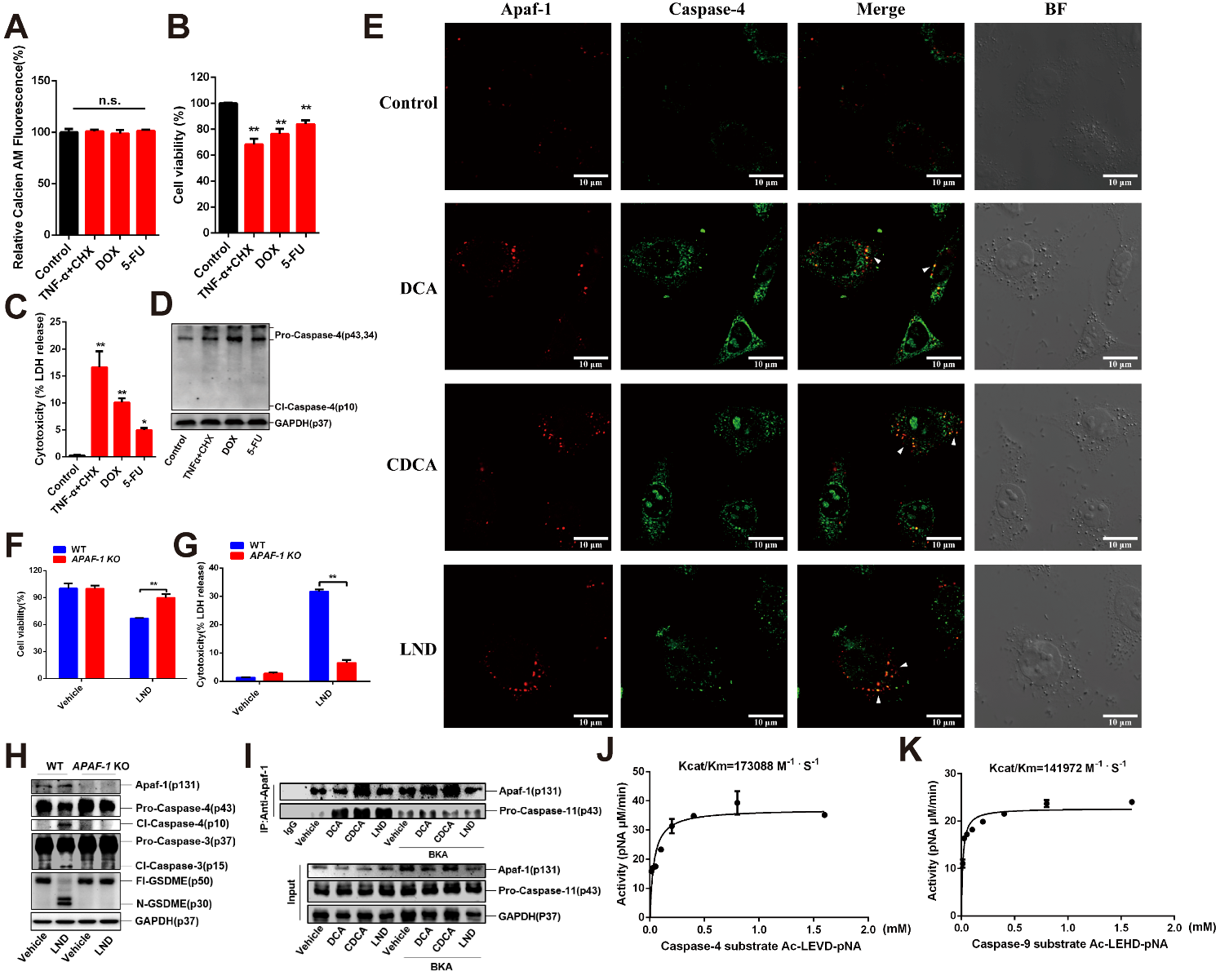

**Figure S5. Both MPT and Apaf-1 Is Necessary for Pyroptosome Assembly. (Related to Figure 5)**

**(A-D)** Calcien-AM/CoCl_2_ assay for MPT **(A)**, CCK-8 assay for cell viability (B), LDH release for cytotoxicity **(C)** and representative immunoblots of caspase-4 and GAPDH in lysates **(D)**. AM/CoCl_2_ assay. HepG2 cells were stimulated with TNFα/CHX (50 ng/mL TNFα + 10 μg/mL CHX), 10 μM Dox or 20 μM 5-FU for 3 hrs in (A) or 12 hrs in (B-D).

**(E)** Representative images by confocal microscopy analysis of the colocalization between Apaf-1 and caspase-4 in HepG2 cells that were stimulated with 200 μM bile acids for 4 hrs or 1 mM LND for 12 hrs. Scale bars, 10 μm.

**(F-H)** CCK-8 assay for cell viability **(F)**, LDH release for cytotoxicity **(G)** and representative immunoblots of Apaf-1, caspase-3, -4, GSDME and GAPDH in lysates **(H)** of *APAF-1* knockout (KO) HepG2 cells compared to wild type (WT) group; cells were treated with 1 mM LND for 12 hrs.

**(I)** Co-immunoprecipitation analysis of Apaf-1 with caspase-11 in BMDMs lysates. BMDMs were pre-treated with 20 μM BKA for 2 hrs followed by 200 μM bile acid for 1 hr or 1 mM LND for 4 hrs.

**(J and K)** The Michaelis-Menten curve and Kcat/Km of caspase-4 or caspase-9, enzymatic activities of caspase-4 and -9 were detected by substrate Ac-LEVD-pNA and Ac-LEHD-pNA respectively. Incubation systems were constructed as Apaf-1: caspase-4=400 nM: 100 nM and Apaf-1: caspase-9=400 nM: 200 nM.

GAPDH was used as loading control in immunoblot analyses of cell lysates. Bar graphs expressed as mean ±SEM (n=3). *, *p*<0.05; **, *p*<0.01; n.s., non-significant versus control or between indicated groups. The Michaelis-Menten curve was fitted and the value of V_max_ and K_m_ was calculated by using Graphpad prism 6.0, K_cat_ =V_max_/[E], E=enzyme concentration.

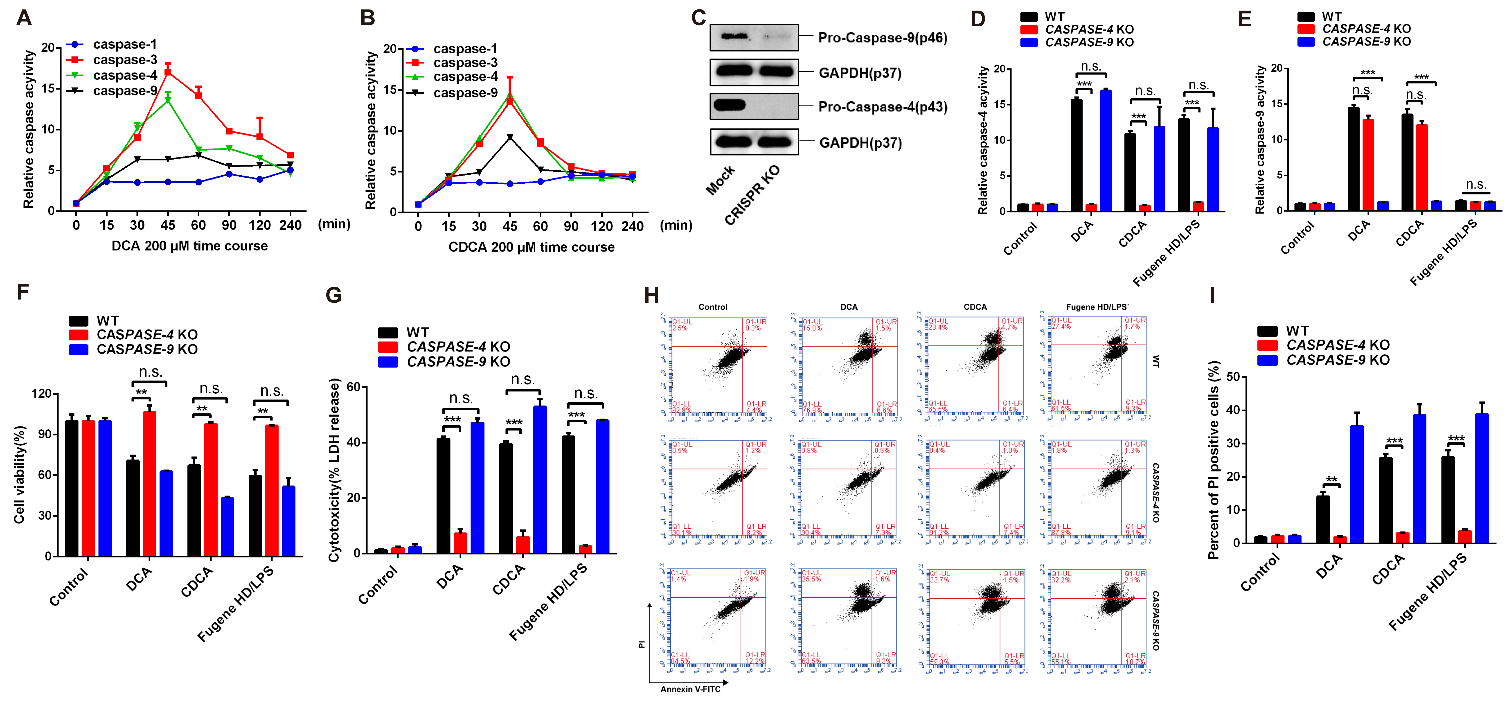

**Figure S6. Caspase-9 Is Not Involved in MPT-Elicited Pyroptosis. (Related to Figure 6)**

**(A and B)** Time course analysis of fold change of caspase-1, -3, -4 and -9 activities in HepG2 cells stimulated with 200 μM bile acid.

**(C)** Immunoblotting validation of CRISPR/Cas9 KO of *CASPASE-4* and *CASPASE-9* efficiency in HepG2 cells.

**(D-G)** Relative caspase-4 and -9 activities (D and E), CCK-8 assay for cell viability **(F)** and LDH release for cytotoxicity **(G)** of *CASPASE-4*/*CASPASE-9* KO HepG2 cells compared to WT group. Cells were treated with 200 μM bile acid for 45 min or LPS-primed HepG2 cells were transfected with 5 μg/ml LPS by 0.3% (v/v) Fugene HD for 10 hrs.

**(H)** Flow cytometry analysis of Annexin V-FITC/PI staining in *CASPASE-4*/*CASPASE-9* KO HepG2 cells compared to WT group, cells were treated as in (D-G).

**(I)** Proportions of PI positive cells, quantified from images as shown in (H).

GAPDH was used as loading control in immunoblot analyses of cell lysates. Line and bar graphs expressed as mean ±SEM (n=3). **, *p*<0.01; ***, *p*<0.001.

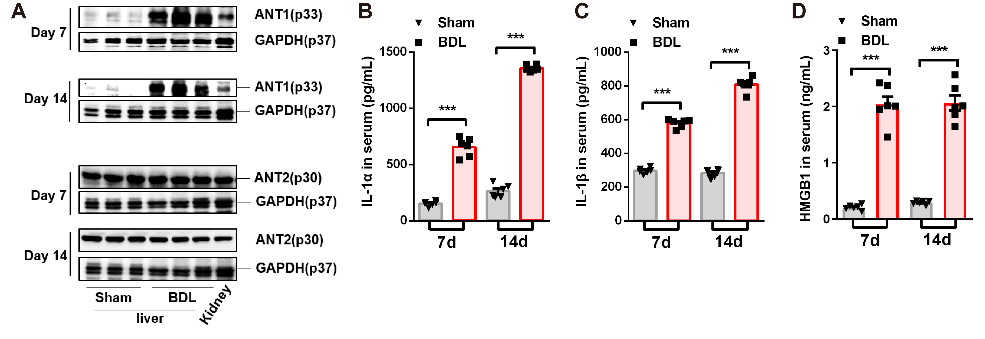

**Figure S7. Caspase-11 and GSDME Dependent Pyroptosis Dominants BDL Induced Cholesteric Liver Failure. (Related to Figure 7)**

**(A)** Representative immunoblots of ANT1/2 and GAPDH in liver and kidney of mice 7 days and 14 days after BDL/sham (n=6).

**(B-D)** Serum levels of IL-1α, IL-1β and HMGB1 in mice 7 days and 14 days after BDL/sham (n=6).

GAPDH was used as loading control in immunoblot analyses of tissue lysates, kidney was analyzed as an indicator of physiological level of ANT, lysates from six mice was merged by two-into-one to n=3. Scatter plot with box graphs expressed as mean ± SEM. ***, *p*<0.001.

**Table S1. Sequences of qPCR Primers. (Related to STAR** **METHODS)**

| **Gene** | **Forward (5’to 3’)** | **Reverse (5’to 3’)** |
| --- | --- | --- |
| ***Homo-CASPASE-4*** | CAGACTCTATGCAAGAGAAGCAACGTATGGCAGGA | CACCTCTGCAGGCCTGGACAATGATGAC |
| ***Mus-Caspase-11*** | ACAAACACCCTGACAAACCAC | CACTGCGTTCAGCATTGTTAAA |
| ***Mus-GAPDH*** | TTGAGGTCAATGAAGGGGTC | TCGTCCCGTAGACAAAATGG |
| ***Homo-GAPDH*** | CCAGGGCTGCTTTTAACTC | GCTCCCCCCTGCAAATGA |
